# GRASS-MIL: graph-based representation and discovery of phenotype-associated spatial structures with multiple instance learning

**DOI:** 10.64898/2026.10.08.757722

**Authors:** Lovro Rabuzin, Josephine Yates, Mina Xu, Valentina Boeva

## Abstract

Tissue function is shaped by both cellular composition and the spatial organization of cells. Spatial single-cell technologies capture this organization across tissue sections, enabling tissue architecture to be linked to disease phenotypes, prognosis and treatment response. However, many current graph-based predictive models depend on assay-specific molecular features, limiting joint analysis of cohorts profiled with incompatible panels, and either aggregate local predictions uniformly or rely on post hoc importance scores that do not directly decompose the basis of a sample-level prediction. Here we introduce GRASS-MIL, a graph multiple-instance learning framework that learns the relative relevance and class-specific evidence of individual cellular neighborhoods, predicts phenotypes across platforms, and decomposes each sample-level prediction exactly into signed contribution from each local neighborhood. These contributions are terms of the prediction itself rather than a post hoc approximation and sum exactly to the predicted logit. In simulations, GRASS-MIL distinguished phenotypes with identical global cell-type composition but different spatial organization, whereas composition-based classifiers performed no better than chance. A single model trained jointly on CosMx spatial molecular imaging, Xenium and MERFISH datasets, harmonized only through a shared cell-type vocabulary, predicted treatment status in high-grade serous ovarian carcinoma with an area under the receiver operating characteristic curve of 0.74 and associated this prediction with two stromal architectures: an immune-rich niche admixed with malignant cells and a desmoplastic niche characterized by malignant-cell exclusion. Applied to spatial proteomics data from non-small-cell lung cancer, GRASS-MIL identified distinct niches associated with relapse (area under the receiver operating characteristic curve, 0.57), including myeloid-rich niches in relapsing tumors and adaptive immune niches in non-relapsing tumors. In a separate lung adenocarcinoma cohort, a similar lymphoid-myeloid polarity was observed when predicting 3-year mortality (area under the receiver operating characteristic curve, 0.62), with a B-cell and CD4+ T-cell niche associated with favorable outcome and a neutrophil and macrophage niche associated with adverse outcome. GRASS-MIL thus combines cross-platform phenotype prediction with an exact, signed decomposition of each prediction into the cellular neighborhoods on which it depends, providing a direct route from sample-level prediction to testable hypotheses linking tissue architecture with patient phenotypes.

## 1 Introduction

Tissue phenotypes are not determined only by which cells are present, but also by how those cells are arranged. In cancer, malignant cells, immune populations, fibroblasts, vasculature and regions of tissue stress form local architectures whose spatial organization is associated with diagnosis, prognosis, treatment response, and immune state [1, 2]. Multiplexed imaging and spatial-omics technologies, including CODEX, imaging mass cytometry, MERFISH, CosMx and Xenium, can measure these architectures at single-cell resolution across large tissue regions [3, 4, 5, 6, 7, 8]. After cell segmentation and cell typing, each sample can be viewed as a spatial graph containing thousands to millions of phenotyped cells. These graphs provide a representation for patient-level prediction from tissue architecture, with the goal of learning which architectures distinguish one phenotype from another. A model’s utility therefore grows with how well it captures local spatial structure, scales to large profiled tissues, operates across assays with different molecular panels, and identifies the tissue structures contributing to their predictions. Many current analyses summarize spatial single-cell data through cell-type abundance, pairwise neighborhood enrichment, cellular-neighborhood clustering or summary point-pattern statistics [9, 10, 11, 12, 13]. These approaches describe tissue composition and organization well, but they typically require hand-chosen scales and do not directly solve weakly supervised, patient-level prediction. In computational pathology, multiple-instance learning (MIL) has become the standard framework for such prediction from whole-slide images, where a slide is represented as a bag of image tiles and only slide-level labels are available [14, 15, 16]; however, tile-based MIL does not model the cellular graph structure that defines spatial single-cell data. Graph-based methods for spatial omics have mostly targeted tasks other than patient-level prediction. One family learns spatial domains or niches without labels by combining expression with spatial adjacency in graph autoencoders, graph convolutions or graph contrastive objectives [17, 18, 19, 20, 21, 22]; another transfers cell-type annotations across datasets while using spatial context [23]; and a third learns general-purpose neighborhood representations, including transformer and foundation models trained across many tissues [12, 24, 25, 26, 27]. These methods produce domain maps, annotations or embeddings that can be provided as input to a downstream classifier, but they do not quantify the contribution of individual neighborhoods to a sample-level prediction, which is the question addressed here. Some recent graph neural network-based approaches, conversely, operate directly on cellular graphs and have been used to predict tissue or disease phenotypes from spatial organization [28, 29, 30, 31].

The three graph-based methods we compare against illustrate the design choices this involves; specifically, what the model takes as input, how a sample-level prediction is formed from many cells, and what kind of interpretive output accompanies it. SPACE-GM [28] is the closest in aim to our work. It extracts a subgraph of fixed radius around every cell, encodes it with a graph network whose node features combine the cell-type label with the measured protein markers, and forms the patient-level prediction by averaging over the resulting subgraph predictions. As a consequence, averaging gives each neighborhood a weight proportional to its abundance rather than to its bearing on the decision. Additionally, the panel-specific marker features prevent cohorts measured with disjoint panels from being analyzed jointly. CytoCommunity [32] instead focuses on supervised niche identification. A graph network with a differentiable graph-partitioning layer assigns cells to tissue cellular neighborhoods, and in its supervised mode the partition is learned jointly with a classifier on sample labels, so that the recovered neighborhoods are those that differ between conditions. Its primary output is a neighborhood partition; the contribution of any one neighborhood to the label of the sample containing it is not a quantity the model defines. SpaCEy [33] takes a third route. It trains a graph network on cell graphs whose node features are the marker abundances themselves, with no cell-type annotation, and interprets the trained model with a separate perturbation-based explainer that learns edge and feature masks under which the prediction is preserved. The mean mask value over a cell’s k-hop neighborhood serves as that cell’s importance, and contiguous runs of high-importance cells mark the informative regions. These importances are magnitudes without sign, so they locate the regions the model relies on but do not say whether a region moves the prediction toward or away from the outcome. Because they come from a second optimization they describe the model rather than decompose it. The simplest spatially informed alternative, which we also evaluate, dispenses with graphs entirely: each sample is summarized by its cell-type composition, either over the whole region or within each local neighborhood, and a standard classifier is trained on these vectors. This family measures how much of a phenotype can be read from abundance alone and therefore provides a composition-based reference against which the added value of spatial structure can be assessed. Across these methods, prediction, attribution and niche discovery are handled by separate components or not at all: SPACE-GM predicts but weights neighborhoods uniformly, CytoCommunity discovers niches but does not score them, and SpaCEy attaches an unsigned post-hoc explainer to its predictor. No existing method combines the three in one model: a patient-level prediction, an exact signed decomposition of that prediction into local neighborhood contributions, and an interpretable organization of those neighborhoods into recurrent tissue motifs.

We introduce GRASS-MIL, a graph multiple-instance learning framework that provides these three outputs within one model. Each sample is represented as a cell graph and treated as a bag of local cellular neighborhoods, which a self-supervised graph neural network encodes and an attention-based MIL head pools into a patient-level prediction. Because the head is additive, each neighborhood receives an attention weight and a class-specific score whose product is its signed contribution to the sample-level class-logit margin, and the margin is exactly the sum of these contributions. The attribution is thus not an approximation of the model’s decision but the decision itself, separated into its parts; grouping neighborhoods by their learned representation then yields the recurrent tissue niches on which the attribution is read out.

We evaluate GRASS-MIL in four settings of increasing complexity, ranging from simulated tissues with known ground truth to real clinical cohorts. A controlled simulation benchmark shows that GRASS-MIL recovers phenotypes defined by mixing, boundary geometry and focal structure, including cases where positive and negative samples share identical cell-type composition and differ only in spatial arrangement, and localizes signed margins to the corresponding ground-truth sources of variation. Applied to high-grade serous ovarian carcinoma (HGSC), a single model trained jointly across CosMx/SMI, Xenium and MERFISH recovers the stromal remodeling known to follow neoadjuvant chemotherapy, resolving it into two architectures: an immune-rich compartment admixed with malignant cells, and a malignant-excluding desmoplastic compartment. Applied to non-small-cell lung cancer (NSCLC), GRASS-MIL predicts relapse and organizes tissue neighborhoods into spatially segregated prognostic territories, with relapse-associated myeloid-rich niches and protective adaptive-immune niches. Applied to lung adenocarcinoma (LUAD), a second NSCLC lung cohort with a different endpoint, GRASS-MIL predicts 3-year mortality and recovers equivalent associations, a protective B-cell and CD4+ T-cell niche opposed by an adverse neutrophil and macrophage niche, whose relative abundance stratifies overall survival. Together, these results indicate that GRASS-MIL combines competitive patient-level prediction with an exact, signed decomposition of each prediction into the cellular neighborhoods it depends on, obtained from the same model that makes the prediction.

## 2 Results

### 2.1 The GRASS-MIL framework to learn interpretable neighborhood representations from cell graphs

GRASS-MIL is a graph-based multiple-instance learning framework that predicts patient-level phenotypes from spatial single-cell data while exposing the local structures behind each prediction. Each sample is represented as a cell graph, with cells as nodes and spatial adjacencies as edges. For every cell, GRASS-MIL extracts the k-hop graph centered on that cell and encodes it with a shared graph neural network (GNN; edge feature-aware GIN layers [28, 34]), yielding one neighborhood embedding per cell. The sample is then treated as a set of these embeddings, which an attention-based MIL head pools into a patient-level prediction (Fig. 1a). The encoder is pretrained without labels using Bootstrapped Graph Latents (BGRL) [35], then fine-tuned end to end on the supervised task (Fig. 1b). Because its input is a harmonized cell-type annotation, optionally with cell size and inter-cell distance, the same framework applies across assays with incompatible marker or gene panels.

**Figure 1.**
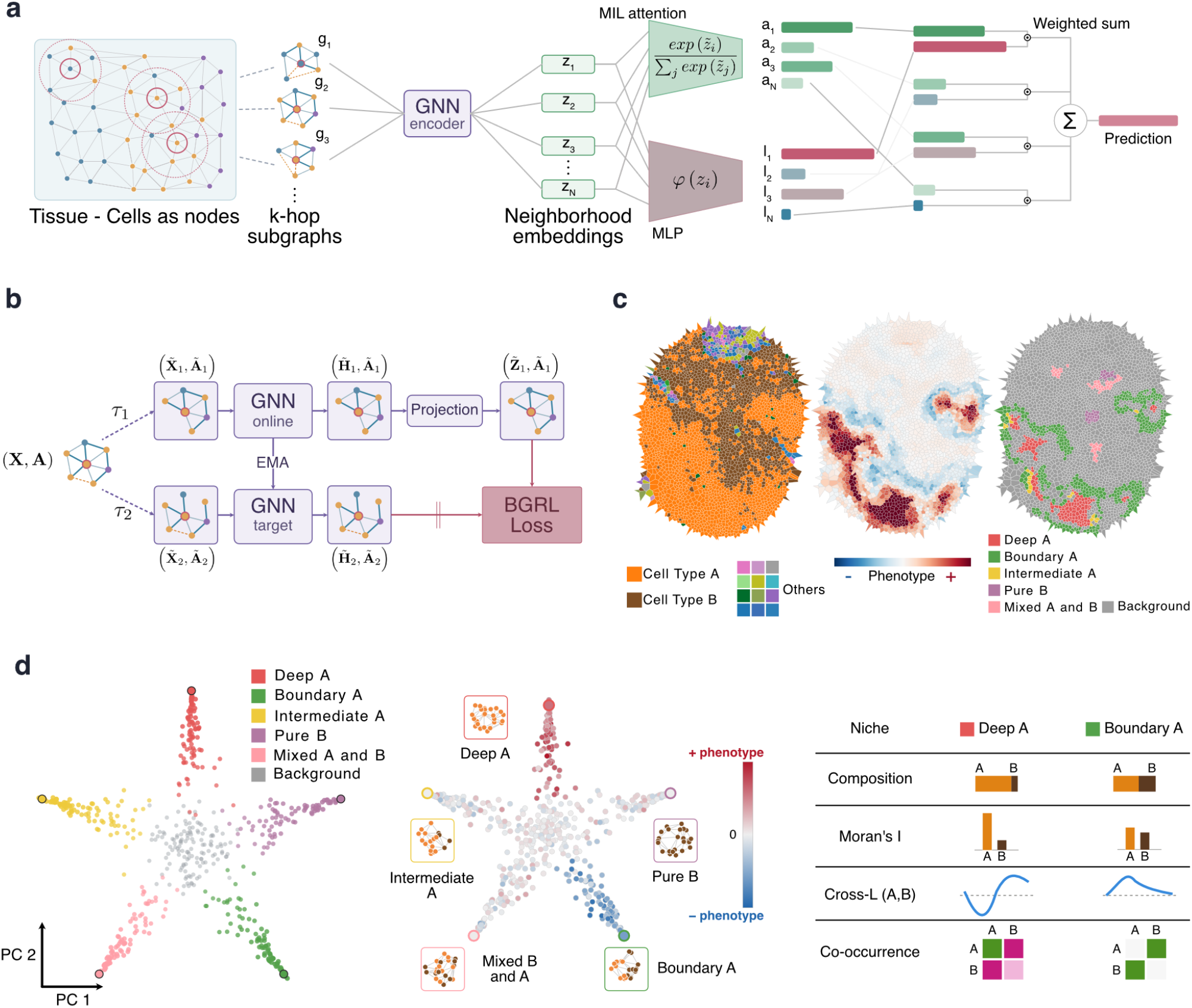
GRASS-MIL predicts patient phenotypes from spatial tissue graphs and resolves an interpretable star-shaped geometry of recurrent tissue niches. **(a)** GRASS-MIL pipeline. Tissue is represented as a cell graph whose nodes are cells carrying cell-type and size features. A k-hop ego-graph is sampled around each cell and mapped by a shared graph neural network (GNN) encoder to a neighborhood embedding *z*_*i*_; the set {*z*_1_, …, *z*_*N*_} contains all neighborhoods of a sample. Attention-based multiple-instance learning (MIL) pools the set through two networks, a gated-attention network assigning each instance a weight 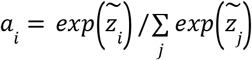, and an MLP network φ producing a per-instance contribution *l*_*i*_. The sample-level prediction is the attention-weighted sum 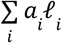 . **(b)** Self-supervised pretraining of the GNN encoder with Bootstrapped Graph Latents (BGRL). Two stochastic augmentations (τ_1_, τ_2_) of the input graph (*X, A*) yield corresponding views 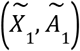 and 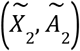. An online encoder and projector are trained to predict the representation of an exponential-moving-average (EMA) target encoder, without negative samples. **(c)** A schematic tissue core (illustrative only) rendered three ways: cells colored by cell type (left), by their contribution to the phenotype prediction (center; diverging scale, negative phenotype in blue, positive phenotype in red), and by recurrent tissue niche (right; six HDBSCAN niches including unassigned ‘Background’). **(d)** Star-shaped geometry and interpretation, shown schematically. Neighborhood embeddings projected onto the first two principal components form a star-shaped structure, with recurrent niches occupying distinct arms of the star and mixed or transitional neighborhoods lying closer to the center (left, colored by HDBSCAN label). The same projection colored by phenotype contribution (center) shows a positive contribution for the Deep A niche and a negative contribution for the A Boundary niche, with the remaining niches near zero. Insets show a representative k-hop ego-graph per niche (nodes colored by cell type: Cell Type A, orange; Cell Type B, brown). Mean cell-type composition per niche (right, top). The centered Ripley’s *L* function (right, bottom) summarizes this second-order spatial structure: relative to the complete-spatial-randomness baseline (CSR), the curve rises to a positive maximum at a characteristic radius ^*^ (co-aggregation above random expectation) before crossing into negative values at larger radii (spatial exclusion), capturing a multi-scale arrangement that cell-type composition alone can not express.

This pooling formulation provides an exact instance-level decomposition of the class-logit margin. Each neighborhood receives an attention weight and a class-margin score, and the sample-level decision is exactly the sum over neighborhoods of their products (Methods). Every neighborhood therefore carries a signed contribution, evidence for or against the predicted phenotype, and these contributions add up to the decision without a baseline, a perturbation or a separately fitted explainer. To make these contributions biologically interpretable, GRASS-MIL clusters the neighborhood embeddings using HDBSCAN [36] with a cosine metric. Dense groups define recurrent tissue niches, while the ‘Background’ label absorbs heterogeneous, transitional or weakly structured neighborhoods rather than forcing them into a discrete niche.

Projected into low-dimensional space, the learned neighborhood embeddings form a star-shaped geometry rather than a diffuse cloud (Fig. 1c,d; confirmed on real cohorts in Fig. 3d and Fig. 4f,g). Recurrent niches occupy distinct arms of the star, while mixed, transitional or weakly structured neighborhoods lie closer to its center and are consequently assigned to the ‘Background’ population. This follows from how density behaves under a cosine metric: recurrent neighborhoods dominated by a single configuration concentrate along one direction, whereas mixtures spread continuously across the directions between them. This geometry is reminiscent of the structure assumed in archetypal analysis, in which observations are represented relative to a small set of extremal configurations [37, 38]. The dense HDBSCAN niches can therefore be interpreted as recurrent configurations near these extremal directions, without specifying their number in advance. Labeling individual embeddings by their signed contribution to the prediction then localizes the phenotype signal to particular arms of the star, identifying which niches contribute positively or negatively to the prediction.

The learned niches capture spatial arrangement beyond cell-type abundance. In the schematic example of Fig. 1d, two niches with near-identical mean cell-type composition occupy different arms of the star and carry different phenotype contributions. As part of its interpretation module, GRASS-MIL profiles each niche with three established spatial statistics that answer complementary questions. Permutation-based neighborhood enrichment [9] describes which cell types sit adjacent to one another more or less often than expected when cell-type labels are shuffled across the same tissue. Moran’s *I* [39] describes whether a cell type forms coherent patches or is scattered cell by cell. Ripley’s *L* and its bivariate cross-*L* functions [40] extend this across distance, describing at what radius cells of one type aggregate around, or are excluded from, cells of another. Together they describe direct contacts, patch coherence and multi-scale arrangement, which are properties that cell-type proportions do not capture.

### 2.2 A controlled simulation benchmark isolates spatial signal beyond composition

To evaluate GRASS-MIL in a setting with known ground-truth spatial organization and cell-type composition, we generated synthetic tissue cores in which the phenotype is determined by spatial arrangement of cell types. Ten phenotype generators span three axes of variation, namely mixing degree, boundary tortuosity and focal-structure presence. The synthetic data deliberately include phenotypes whose positive and negative classes share identical global cell-type composition (Fig. 2a,b). Boundary-proximal neighborhoods were excluded from evaluation for all methods, to avoid finite-window artifacts (Methods).

**Figure 2.**
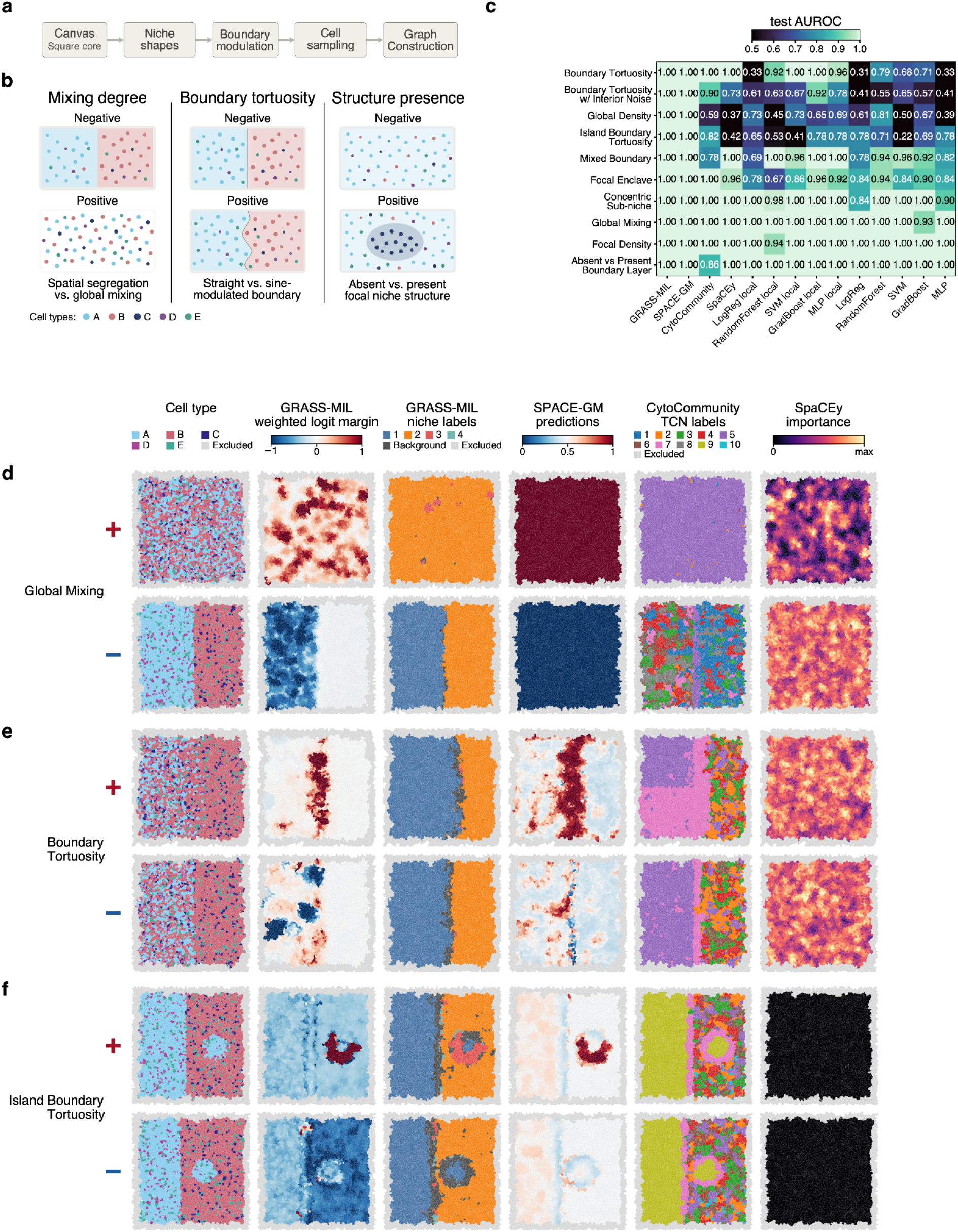
A controlled simulation benchmark shows that GRASS-MIL recovers spatial phenotypes spanning mixing, boundary geometry and focal structure, including arrangements that composition-based classifiers cannot detect. **(a)** Generative pipeline. Each synthetic core is built on a square canvas; niche compartments are laid out as rectangles or ellipses, their boundaries are optionally modulated by a sine function, cells are sampled by a uniform Poisson process and assigned types, and a Delaunay triangulation yields the cell graph passed to all methods. **(b)** Three controlled axes of variation, each contrasting a negative and a positive phenotype: mixing degree, boundary tortuosity and focal-structure presence. Cell types A to E are color-coded throughout. **(c)** Per-phenotype test AUROC across ten phenotype generators and fourteen methods, evaluated after excluding boundary-proximal neighborhoods. Methods comprise GRASS-MIL, three spatial-niche baselines (SPACE-GM, CytoCommunity, SpaCEy) and standard classifiers in two variants: a global variant trained on whole-sample cell-type composition and a local variant trained on individual 4-hop subgraph compositions with mean aggregation to the sample level. The comparison separates the effect of local aggregation from the effect of learned graph representations. **(d–f)** Representative simulated cores for Global Mixing (d), Boundary Tortuosity (e) and Island Boundary Tortuosity (f), each shown for the negative (top) and positive (bottom) phenotype under six renderings: cell type, with excluded boundary-proximal cells marked ‘Excluded’; GRASS-MIL attention-weighted logit margin; GRASS-MIL niche assignment; SPACE-GM per-cell prediction; CytoCommunity tissue-cellular-neighborhood (TCN) label; and SpaCEy node importance. In (f), SpaCEy importance is identically zero for both cores because its classifier collapsed to a constant prediction on this generator. GRASS-MIL margins localize to the discriminative spatial structures that distinguish positive from negative cores.

GRASS-MIL was compared with thirteen baselines: three spatial-niche methods, SPACE-GM [28], CytoCommunity [32] and SpaCEy [33], and five standard classifiers in two feature settings: global and local, differing in the scale at which composition is measured. The global setting represents each sample by a whole-sample cell-type composition vector. The local setting represents each 4-hop subgraph by its local cell-type composition, predicts at the subgraph level and mean-aggregates predictions to the sample level. All methods, including SPACE-GM and SpaCEy, received the same cell-type input as GRASS-MIL, with cell size where available, to keep the input classes comparable across methods so that the comparison isolates the modeling approach rather than the input. Results for SPACE-GM and SpaCEy trained on their native marker-intensity inputs are reported in Supplementary Fig. 1; with input classes matched, GRASS-MIL achieved the highest mean AUROC among the evaluated methods. Both GRASS-MIL and SPACE-GM achieved perfect discrimination across all simulated phenotype generators (Fig. 2c). The other two spatial-niche methods achieved near-perfect performance on most generators but not all: CytoCommunity dropped to 0.59 on Global Density, and SpaCEy fell below chance on Global Density (0.37) and Island Boundary Tortuosity (0.42, Fig. 2c,f). Composition-based classifiers were less robust, especially on phenotypes for which positive and negative samples share identical global composition and differ primarily in spatial arrangement, including Boundary Tortuosity and Global Density (Fig. 2c). Local composition features improved over global composition on average, but did not close the gap to learned graph representations. Thus, the simulation benchmark separates three levels of information: global cell-type abundance, local cell-type abundance and graph-encoded spatial arrangement.

The signed GRASS-MIL margins localized to the structures that define the simulated phenotypes. In held-out examples from Global Mixing, Boundary Tortuosity and Island Boundary Tortuosity, high-magnitude margins appeared around the discriminative spatial motifs rather than being distributed uniformly across the core (Fig. 2d–f; the remaining generators are shown in Supplementary Fig. 3). Because the generative spatial structures are known, the simulation provides a direct setting in which to assess whether the model’s neighborhood contributions localize to the structures defining the phenotype. The model’s own decision, split into neighborhoods, is consistent with the ground-truth construction of the benchmark and supports the use of per-neighborhood margins for downstream niche interpretation. The baselines’ readouts behaved differently on the same cores: SPACE-GM’s per-cell predictions also highlighted the discriminative motifs, CytoCommunity’s TCN labels partitioned the cores into compartments without attaching a direction to any of them, and SpaCEy’s node importances were spread across the core in the Global Mixing and Boundary Tortuosity examples and were identically zero for Island Boundary Tortuosity, the generator on which its classifier collapsed (Fig. 2d–f). Together, these results show that GRASS-MIL recovers spatial phenotypes that cell-type composition alone cannot express, and that its per-neighborhood margins localize to the structures used to define the simulated phenotype.

### 2.3 GRASS-MIL resolves a cross-platform stromal-architecture axis predicting treatment status in HGSC

To evaluate the performance of GRASS-MIL in a real-world application, we next applied it to high-grade serous ovarian carcinoma (HGSC) using the spatial atlas of Yeh et al. [41] to predict treatment status. This atlas comprises five acquisitions across three spatial-omics platforms, CosMx/SMI, Xenium and MERFISH, spanning 94 patients and disjoint gene panels with profiled gene counts ranging from 140 to 6,175 (Fig. 3a). To train a joint model predicting treatment status across platforms, all datasets were reduced to a shared vocabulary of ten cell types plus ‘Other’. Cell-type harmonization was therefore the only integration step required: because the model uses cell-type labels and spatial geometry alone, the five acquisitions could be pooled without a common gene space, batch correction or expression-level integration. A molecular-level model would instead require additional cross-platform harmonization to accommodate the disjoint gene panels. Under this harmonized representation, GRASS-MIL achieved the highest mean test AUROC for treatment-status prediction among the evaluated methods under nested 5-fold cross-validation (AUROC = 0.74 ± 0.06), exceeding the best competing spatial-niche method (SpaCEy, 0.71) and the best composition-based classifier (0.72) by two to three percentage points (Fig. 3g; Supplementary Table 3).

**Figure 3.**
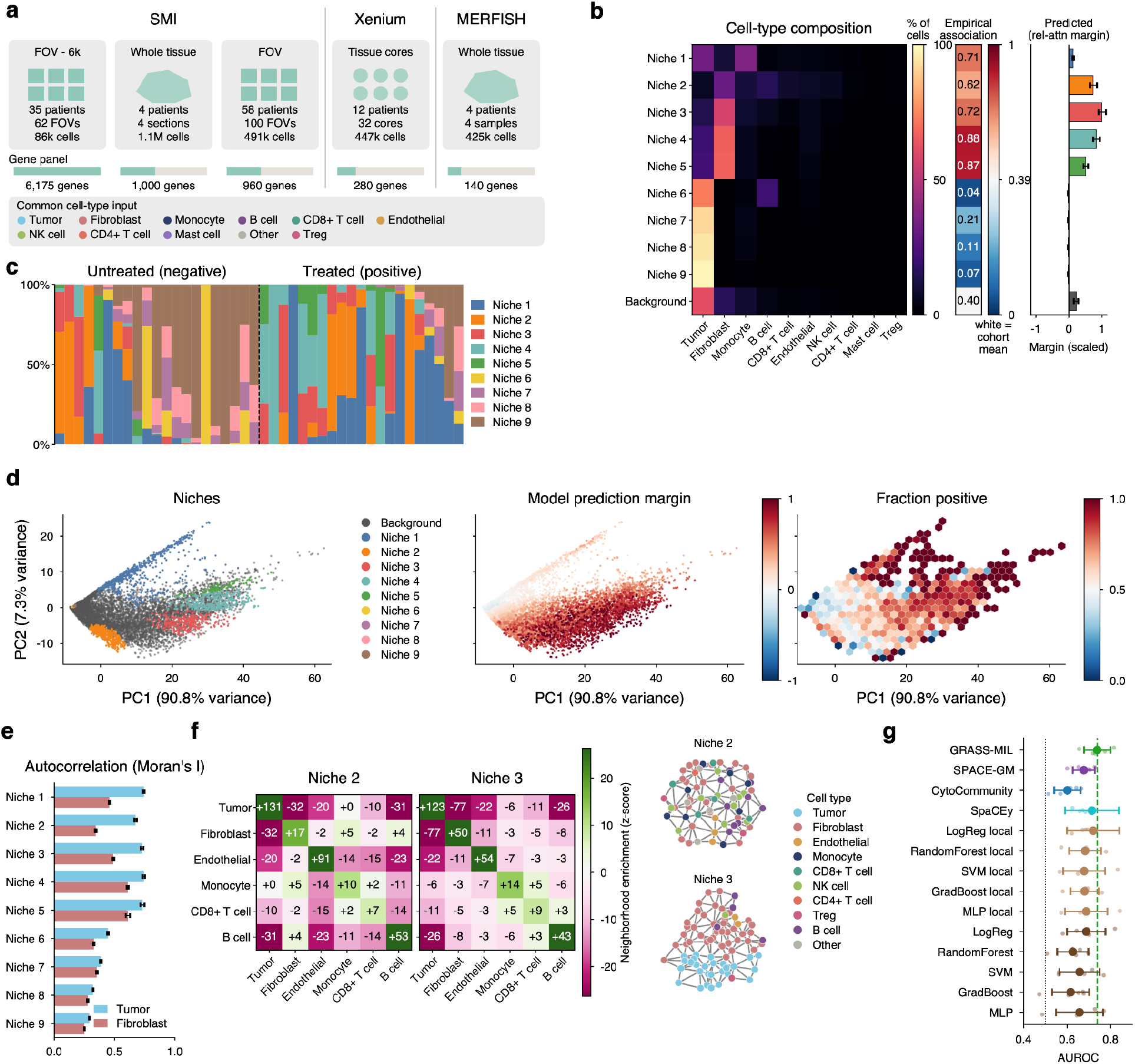
GRASS-MIL resolves a stromal-architecture axis that predicts treatment status in high-grade serous ovarian carcinoma (HGSC) and transfers across spatial platforms. **(a)** Five HGSC datasets across three platforms (CosMx/SMI, Xenium, MERFISH; 94 patients, ∼2.5M cells, disjoint 140-6,175-gene panels) are reduced to a shared vocabulary of ten cell types plus ‘Other’. Only cell-type labels enter the model, making GRASS-MIL assay-agnostic after cell-type harmonization. **(b)** Instances are clustered with HDBSCAN in the gated-attention head space into nine niches and ‘Background’. Left to right: mean cell-type composition, empirical treated fraction and relative-attention margin. Attribution concentrates in stromal niches 2-5, with peak positive margin in niche 3, whereas niches 6-9 carry margins near zero. **(c)** Per-sample niche composition by treatment status. Treated samples are enriched for high-attribution stromal niches. **(d)** PCA of the instance embedding (PC1, 90.8%; PC2, 7.3%) colored by niche, prediction margin and empirical treated fraction. The margin varies continuously along PC1. **(e)** Per-niche Moran’s *I* for malignant and fibroblast cells (mean ± s.e.m. across instances), indicating spatial clustering. Stromal niches 2-5 are more spatially coherent than near-null niches 6-9; niche 3 exemplifies a high-coherence desmoplastic regime. **(f)** Neighborhood enrichment (squidpy permutation *z*-scores) with representative Delaunay graphs for two top-attributed niches. Both correspond to post-NACT desmoplastic stroma but differ architecturally: niche 2 is immune-rich and admixed with malignant cells, whereas niche 3 is a tumor-stroma boundary with coherent fibroblast sheets and malignant-cell exclusion. (g) Test AUROC under nested 5-fold cross-validation (dots, folds; marker and bar, mean ± s.d.; dashed line, GRASS-MIL; dotted line, chance).

HDBSCAN clustering of HGSC neighborhood instances identified nine recurrent niches and the unassigned Background population. Most positive attribution for the treated-versus-untreated decision concentrated in a contiguous set of stromal niches, niches 2-5, with the largest contribution in niche 3 (Fig. 3b). Niches 6-9 carried margins near zero, contributing little in either direction to the decision. Treated samples were enriched for the high-attribution stromal niches (Fig. 3b,c). The empirical treated fraction and the attribution did not always agree, however. Niche 1, a tumor and monocyte niche, had an empirical treated fraction of 0.71 against a cohort mean of 0.39 but a small margin, and the tumor-dominated niches 6-9, whose cells came almost entirely from untreated sections (empirical fractions 0.04-0.21), carried no attribution. The empirical fraction is a marginal, cell-weighted statistic that large sections dominate, whereas the attribution is the contribution of a niche to a prediction made with every other neighborhood of the sample present; a niche that is prevalent in one class but redundant given the stromal niches receives little weight (Discussion). In the instance embedding, the prediction margin varied smoothly along the leading principal component (PC1, 90.8% variance explained), indicating that treatment status is represented primarily along a continuous stromal-architecture axis rather than by isolated rare motifs (Fig. 3d).

Per-niche Moran’s I indicated that the high-attribution niches differ from the near-zero ones not only in stromal content but in how that stroma is organized. Fibroblasts and malignant cells were more spatially autocorrelated, meaning grouped into coherent patches rather than scattered cell by cell, in niches 2-5 than in niches 6-9 (Fig. 3e). Therefore the treatment-associated niches are not merely fibroblast-enriched but contain spatially structured fibroblast and tumor compartments. This distinction matters because post-treatment stroma of similar abundance can be organized in architecturally distinct ways, which measures of stromal content alone cannot separate.

Neighborhood-enrichment analysis further separated the two highest-attribution niches into distinct stromal architectures (Fig. 3f). For each niche we permuted cell-type labels over the fixed spatial graph and scored how often each pair of cell types was adjacent relative to this null, reporting the result as a z-score, where positive values indicate co-localization and negative values spatial exclusion. Niche 2 was an immune-rich desmoplastic compartment: B cells were strongly co-localized with one another (z = +53), monocytes and CD8+ T cells were associated, and malignant cells remained admixed throughout, consistent with a fibro-inflammatory treatment-associated environment in which stromal remodeling and immune infiltration co-occur. Niche 3 instead represented a sharper tumor–stroma boundary, with fibroblasts forming coherent sheets (z = +50) and rarely adjacent to malignant cells (z = −77). Interestingly, both niches showed similarly high fibroblast Moran’s I, despite having very different spatial organizations. In Niche 3, fibroblasts were strongly excluded from malignant cells and formed a continuous border around the tumor, consistent with the exclusion of CD8+ T cells by cancer-associated fibroblasts described in solid tumors [42]. In Niche 2, by contrast, fibroblasts occurred in smaller clusters interspersed with other cell types. Thus, similar spatial statistics can capture very different tissue architectures. By learning the underlying spatial organization, our approach can distinguish biologically distinct niches that would otherwise appear similar based on conventional spatial metrics alone.

These two niches provide complementary spatial interpretations of the treated state, an immune-admixed stromal response (niche 2) and a malignant-excluding desmoplastic boundary (niche 3), and together they are consistent with the fibro-inflammatory and desmoplastic stromal changes reported after neoadjuvant chemotherapy [43, 44] (Discussion).

This treatment-status signal is also distinct from the principal finding of the source atlas [41], which focused on a malignant cell state associated with T- and NK-cell infiltration and checkpoint-blockade response as opposed to investigating patterns associated with treatment status. In the task analyzed here, malignant niches carry little attribution, whereas the discriminative signal resides primarily in stromal architecture. GRASS-MIL therefore recovers a task-specific spatial correlate of treatment exposure: organized fibro-inflammatory and desmoplastic tissue remodeling across platforms.

### 2.4 GRASS-MIL predicts NSCLC relapse and organizes neighborhoods into spatially segregated prognostic territories

As a second real-world application, GRASS-MIL was applied to relapse prediction in a large non-small-cell lung cancer (NSCLC) imaging mass cytometry cohort comprising 1,070 patients and 2,070 tissue microarray cores [45, 46], of which 1,868 cores were retained after restricting the cohort to adenocarcinoma, squamous-cell and adenosquamous histology (Methods). We used the published annotation of 30 cell types spanning tumor, cancer-associated fibroblast (CAF), lymphoid and myeloid lineages [45] (Fig. 4a). Because all samples were acquired on one platform under a common segmentation, cells could additionally carry segmentation-derived cell area, providing an additional morphology-related feature. This prediction task presents additional challenges: reported associations between NSCLC microenvironmental organization and outcome vary by histology and by the maturity and location of the structures involved [47, 48, 49]. Moreover, each core samples approximately 1 mm^2^ of the tumor leading edge, introducing uncertainty as to whether relevant spatial structures are represented in a given core. Consistent with these challenges, none of the evaluated classifiers achieved strong predictive performance. GRASS-MIL achieved the highest mean test AUROC for relapse prediction among the evaluated methods under nested 5-fold cross-validation (AUROC = 0.57 ± 0.01). This exceeded the best competing spatial-niche method by 4 percentage points (SPACE-GM, 0.53) and the best composition-based classifier by 1.5 percentage points (GradBoost local, 0.56; Fig. 4h; Supplementary Table 3).

**Figure 4.**
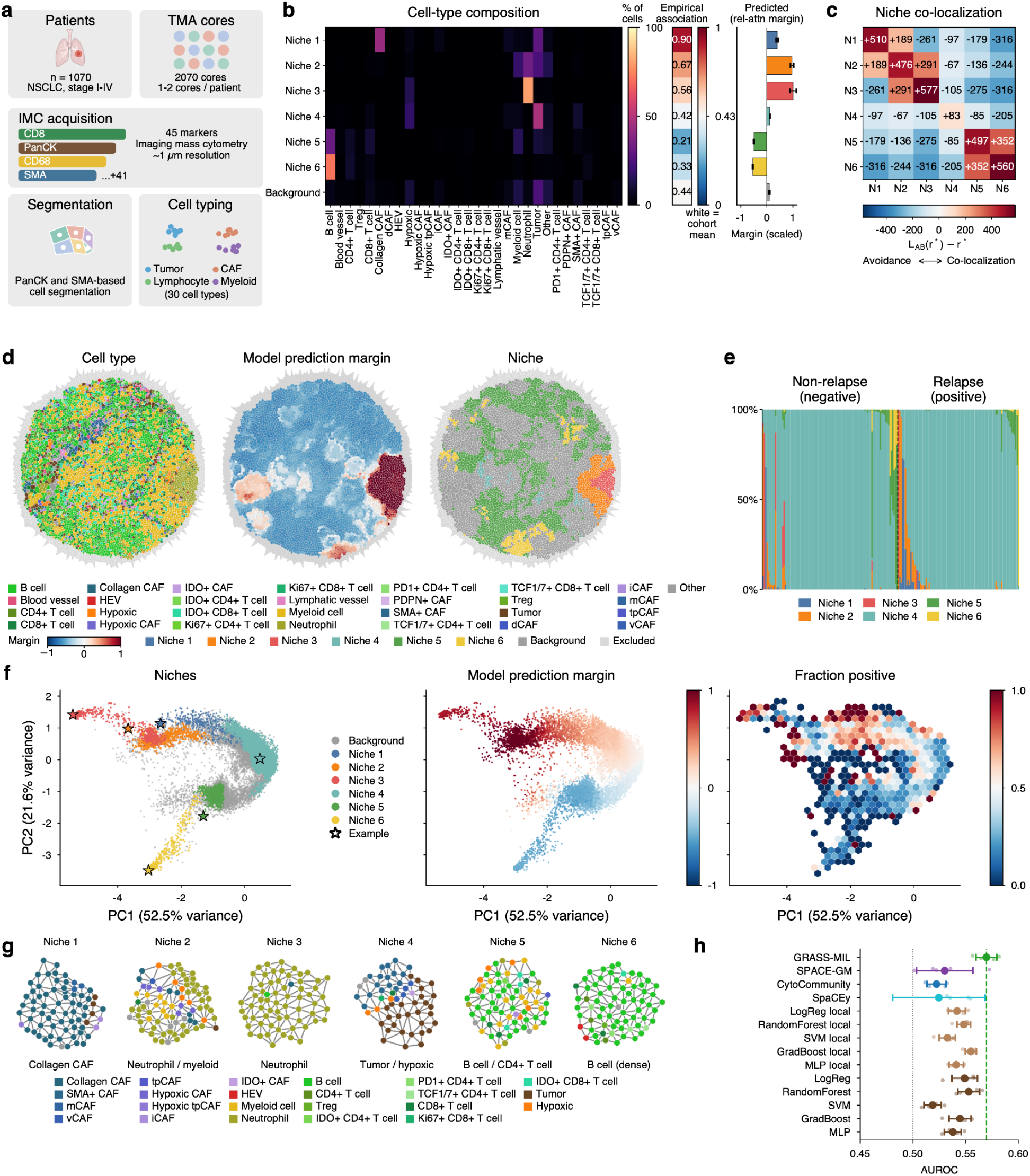
GRASS-MIL predicts relapse from NSCLC tissue architecture and organizes cellular neighborhoods into spatially segregated, prognostically opposed niches. **(a)** Cohort and pipeline. An NSCLC imaging mass cytometry cohort (1,070 patients; 2,070 TMA cores, one to two per patient; 45 markers, ∼1 µm resolution) is segmented and typed into 30 cell types spanning tumor, CAF, lymphoid and myeloid lineages. The task is relapse prediction. **(b)** Niche definition and attribution. Instances are clustered with HDBSCAN in the gated-attention head space into six niches plus ‘Background’. Left to right: mean cell-type composition, empirical relapse fraction and per-niche relative-attention margin. Niches 1-3 carry positive, relapse-associated margins (largest for niches 2-3), whereas niches 5-6 carry negative, protective margins. **(c)** Meso-scale niche organization quantified by centered Ripley’s cross-*L* at the characteristic radius *r*^*^, *L*_*AB*_ (*r*^*^ ) − *r*^*^ (positive, co-aggregation; negative, spatial exclusion; diagonal, within-niche self-aggregation). Niches partition into two spatial territories: a relapse-associated chain in which niche 2 co-aggregates with niches 1 and 3, which avoid each other, and a separate co-aggregating block containing niches 5-6. Niche 4 is weakly self-aggregating and spatially diffuse. **(d)** Representative TMA core rendered by cell type, per-cell model margin and niche assignment. **(e)** Per-sample niche composition by relapse status. Relapse samples are enriched for niches 1-3 and depleted of protective niches 5-6. **(f)** PCA of the instance embedding (PC1, 52.5%; PC2, 21.6%) colored by niche, prediction margin and empirical relapse fraction, with Background at the center and niches at the periphery; niches 1-3 occupy distinct segments of one arc, niches 5-6 distinct segments of a second arc, and niche 4 lies closer to the center. Stars mark ego-graphs shown in (g). **(g)** Representative 4-hop subgraphs for each niche, with nodes colored by cell type. (h) Test AUROC under nested 5-fold cross-validation (dots, folds; marker and bar, mean ± s.d.; dashed line, GRASS-MIL; dotted line, chance).

HDBSCAN clustering of NSCLC instances in the gated-attention head space identified six recurrent niches plus an unassigned Background population (Fig. 4b). The niches fell into three compositional groups. Niches 1–3 were stromal and myeloid: niche 1 was dominated by collagen CAFs with admixed tumor cells and, after column-wise z-scoring that exposes rarer CAF subtypes, by hypoxic, tumor-promoting and matrix CAFs (Supplementary Fig. 2b); niche 2 combined neutrophils, myeloid cells and tumor cells; and niche 3 consisted almost entirely of neutrophils. Niche 4 was a tumor-dominated niche with hypoxic cells. Niches 5–6 were lymphoid: niche 5 mixed B cells with CD4+ T cells, whereas niche 6 was a dense B-cell aggregate. Attribution separated these groups by prognostic sign. Niches 2 and 3 carried the largest positive, relapse-associated margins; niche 1 was also relapse-associated, with the highest empirical relapse fraction of any niche (0.90 versus a cohort mean of 0.43) but a smaller margin; niches 5 and 6 carried negative margins (empirical relapse fractions 0.21 and 0.33); and niche 4 and Background lay near zero (Fig. 4b). In a representative core, the margin field tracked niche assignment, with positive margins over the neutrophil-rich territory of niches 2-3 and negative margins over the lymphoid niches 5-6 (Fig. 4d). Per-sample niche composition showed the same split, with relapse samples enriched for niches 1-3 and depleted of niches 5-6 (Fig. 4e). Relapsing samples therefore contained a greater proportion of neutrophil- and myeloid-rich niches and a lower proportion of lymphoid, TLS-like niches.

The spatial organization of these niches showed a compartmentalized tissue architecture. To quantify it, we used Centered Ripley’s cross-*L* at the characteristic radius, *L*_*AB*_ (*r*^*^) − *r*^*^. Intuitively, this statistic asks, for each pair of niches, how many neighborhoods of one niche lie within the characteristic radius of a neighborhood of the other, relative to the number expected if the two niches were distributed independently at their observed densities: it is zero under independence, positive when the two niches co-aggregate and negative when they avoid each other, with the characteristic radius chosen as the distance at which these departures are largest across all niche pairs (Methods). Centered Ripley’s cross-*L* partitioned the niches into relapse-associated and protective territories that exclude each other (Fig. 4c). Niche 2 co-aggregated with both niche 1 and niche 3, which in turn avoided each other, so the relapse-associated niches formed a chain of adjacent territories rather than one mixed compartment; niches 5-6 formed a separate co-aggregating block that excluded all three relapse-associated niches. The relapse-associated and protective niches therefore occupied distinct tissue territories rather than being uniformly intermingled. Niche 4 showed weak self-aggregation and near-null attribution, consistent with a more spatially diffuse, less predictive population.

The NSCLC instance embedding again placed Background at the center and the niches at the periphery, but the relapse-associated niches 3, 2 and 1 occupied distinct segments of one arc, in the same order as their spatial adjacency, the protective niches 5 and 6 occupied distinct segments of a second arc, and the near-null niche 4 lay closer to the center (Fig. 4f; representative ego-graphs for each niche in Fig. 4g). This geometry matters for interpretation because the model margin varied smoothly from near zero at the center to its extreme at the far end of each arc, and the fraction of neighborhoods drawn from relapsing cores tracked the margin along the same paths (Fig. 4f, middle and right). Niches are therefore the endpoints of continuous transitions through intermediate neighborhoods rather than isolated islands, and attribution agreed with outcome along the full transition, not only at the niche level. Across the synthetic benchmark, HGSC and NSCLC, the learned neighborhood space thus arranged tissue into recurrent structural motifs at its periphery, and in NSCLC the arrangement of those motifs in the embedding reflected their adjacency in tissue.

Niche 1 recapitulated previously reported associations between CAF phenotypes and prognosis in this cohort. Cords et al. defined 11 CAF phenotypes in these data and found collagen, hypoxic, tumor-like and matrix CAFs to be associated with shorter survival and with immune-excluded microenvironments, whereas SMA+ CAFs, inflammatory CAFs and interferon-response CAFs marked inflamed microenvironments and longer survival [45]. Column-wise z-scoring of the niche compositions, which exposes CAF subtypes that are rare in absolute terms (Supplementary Fig. 2b), shows that niche 1 was enriched for these poor-outcome-associated CAF phenotypes, most strongly for collagen CAFs, followed by hypoxic tumor-promoting CAFs, hypoxic CAFs and matrix CAFs, and depleted of SMA+ CAFs, PDPN+ CAFs, vascular CAFs and every immune population; inflammatory CAFs were the only CAF phenotype previously associated with favorable prognosis that was also enriched. Niche 1 accordingly had the highest empirical relapse fraction of any niche (0.90), and its immune depletion matches the reduced immune infiltration that Cords et al. reported for matrix-CAF-high tumors [45]. GRASS-MIL therefore recovers, from cell-type labels and geometry without marker intensities, the CAF phenotypes and the immune exclusion that the original marker-level analysis of this cohort associated with poor prognosis.

By contrast, the myeloid niches represent a spatial pattern not isolated in the original analysis. The largest relapse attribution fell on the neutrophil-rich niches 2 and 3, which are compositionally distinct from niche 1 and spatially arranged as a chain, with niche 2 co-aggregating with both niche 1 and niche 3 while niches 1 and 3 avoid each other (Fig. 4b,c). In the CAF-centered analysis of Cords et al., neutrophils appeared only as one of several cell types enriched in the poor-prognosis patient groups [45]; here they define the neighborhoods that carry most of the relapse signal, spatially segregated from the protective lymphoid niches. This myeloid axis is consistent with independent reports linking neutrophils to recurrence and poor outcome in NSCLC, whether measured as the intratumoral neutrophil-to-CD8 ratio [50] or as the systemic neutrophil-to-lymphocyte ratio [51], with the caveat that the prognostic effect of intratumoral neutrophils can depend on histology [49]. The protective niches 5–6 are adaptive-immune and TLS-like, consistent with reports linking mature or intratumoral TLSs to favorable NSCLC outcomes, while TLS prognostic value can vary with maturity and spatial location [47, 48, 52].

Niche 1 contained the matrix-dense and hypoxic CAF phenotypes that Cords et al. associated with immune exclusion [45]. The niches bordering it were enriched for hypoxic tumor cells and neutrophils (Supplementary Fig. 2b), yet attribution peaked in the neutrophil-dense niche 3 rather than in the stroma itself. Hypoxia alone did not carry the signal: niche 4, which carries the greatest share of hypoxic tumor cells, had a margin near zero. The relapse signal therefore localized to neutrophil-dense tissue adjacent to hypoxic stroma, rather than to the stroma or to hypoxic tumor cells themselves (Discussion).

### 2.5 GRASS-MIL predicts 3-year mortality in lung adenocarcinoma from a lymphoid–myeloid niche axis

To test whether the lung findings generalize to a second cohort and a different endpoint, we applied GRASS-MIL to the lung adenocarcinoma (LUAD) imaging mass cytometry cohort of Sorin et al. [53], in which one 1-mm^2^ core per patient was profiled with a 35-marker panel and resolved into 17 cell types (Fig. 5a). The task was to predict death within 3 years of surgery; after excluding patients censored before 3 years, 407 patients with 89 deaths remained. On this task, GRASS-MIL achieved the highest mean test AUROC among the evaluated methods under nested 5-fold cross-validation (AUROC = 0.62 ± 0.07), exceeding SPACE-GM (0.58), SpaCEy (0.57) and the best composition-based classifier (0.59; Fig. 5f; Supplementary Table 3).

**Figure 5.**
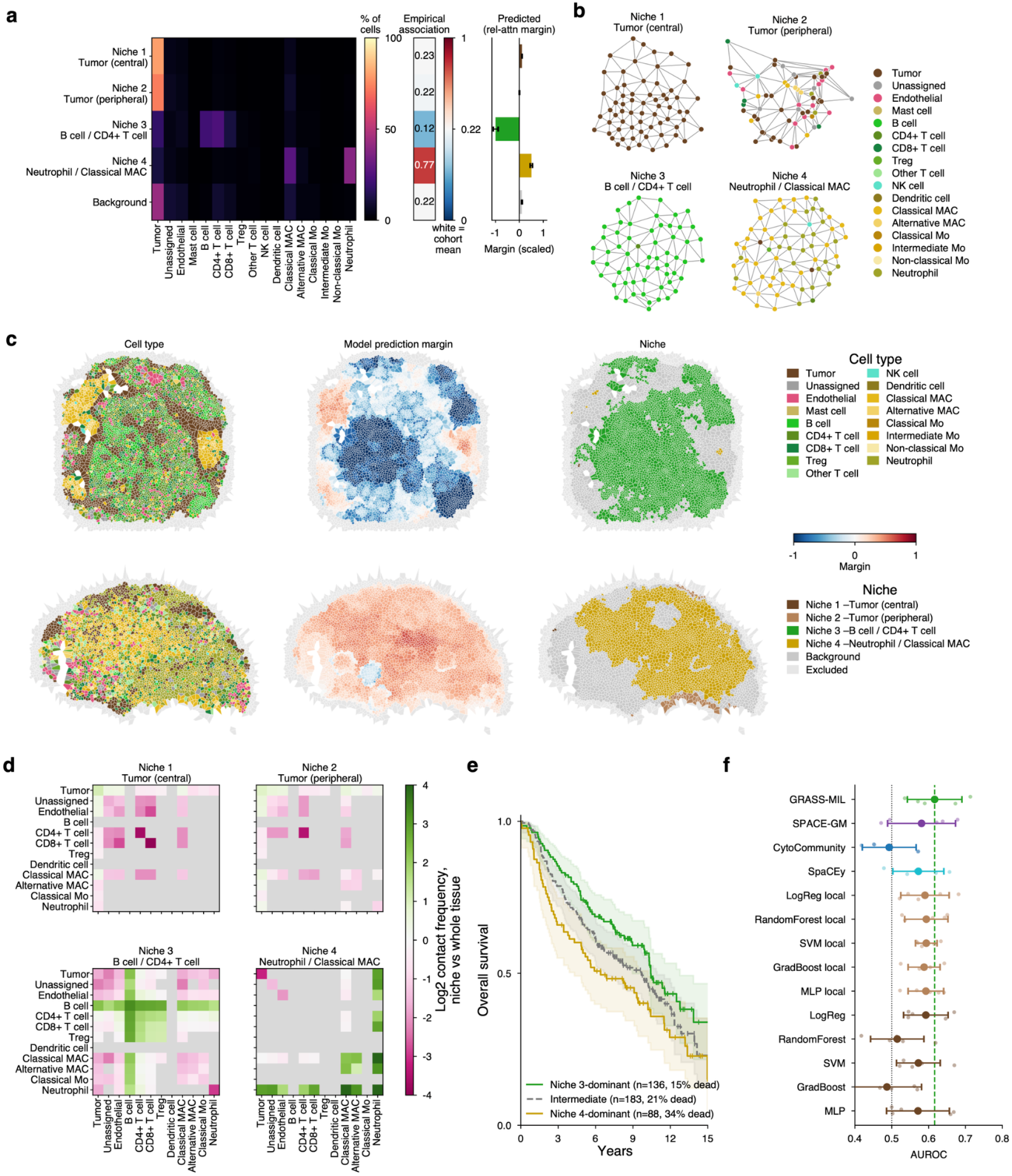
GRASS-MIL predicts 3-year mortality from LUAD tissue architecture and resolves a protective lymphoid niche and an adverse myeloid niche. **(a)** Niche definition and attribution. A lung adenocarcinoma imaging mass cytometry cohort (407 TMA cores, one per patient, 17 cell types) is used to predict death within 3 years of surgery; instances are clustered with HDBSCAN in the embedding space into four niches plus ‘Background’ and every cell of every core is assigned by nearest neighbor. Left to right: cell-type composition (% of niche cells), per-niche relative-attention margin with bootstrap confidence intervals (positive values indicate contributions toward the death-within-3-years class), fraction of niche cells from patients who died within 3 years (dashed line, cohort average) and niche share. Niches 1-2 are central and peripheral tumor niches with neutral margins, niche 3 (B cell / CD4+ T cell) carries a negative, protective margin and niche 4 (neutrophil / classical macrophage) a positive, adverse margin. **(b)** Representative 4-hop subgraphs for each niche, with nodes colored by cell type. **(c)** Two held-out TMA cores rendered by niche assignment, cell type and per-cell model margin; the 4-hop rim excluded from the model is shown in grey. **(d)** Within-niche cell-type contact enrichment, log2 contact frequency inside the niche relative to the whole tissue (grey, fewer than 100 contacts). Niche 3 is organized around B-cell-B-cell and B-cell-T-cell contacts, niche 4 around neutrophil and macrophage contacts. **(e)** Kaplan-Meier overall survival for all 407 patients stratified into tertiles of a two-niche score, the z-scored log share of niche 3 minus that of niche 4 (three-group log-rank p = 0.020; Cox hazard ratio 0.89 per unit, p = 0.003). Niche 3-dominant patients have the best and niche 4-dominant patients the worst survival. **(f)** Test AUROC under nested 5-fold cross-validation (dots, folds; marker and bar, mean ± s.d.; dashed line, GRASS-MIL; dotted line, chance). GRASS-MIL achieved the highest mean AUROC among the evaluated methods.

HDBSCAN clustering of neighborhoods identified four recurrent niches plus Background (Fig. 5a,b). Niches 1 and 2 were tumor niches that differed in position rather than in lineage: niche 1 captured the central tumor, with subgraphs composed almost entirely of tumor cells, whereas niche 2 captured the tumor periphery, where tumor cells were admixed with endothelial, mast and immune cells. Niche 3 was lymphoid, combining B cells with CD4+ T cells, and niche 4 was myeloid, combining neutrophils with classical macrophages. Attribution separated these immune niches by prognostic sign while leaving the tumor niches neutral. Niche 3 carried a strong negative, protective margin, with 0.12 of its cells from patients who died within 3 years against a cohort mean of 0.22, whereas niche 4 carried a positive, adverse margin and a fraction of 0.77; niches 1–2 and Background lay near zero, with fractions close to the cohort mean (Fig. 5a). The same separation was visible within individual held-out cores: a core dominated by niche 3 received negative margins throughout its lymphoid aggregates, whereas a core dominated by niche 4 received positive margins throughout its neutrophil- and macrophage-rich territory (Fig. 5c).

Within-niche contact enrichment showed that the two prognostic niches were built from distinct cell–cell contacts rather than merely from distinct cell counts (Fig. 5d). Relative to the whole tissue, niche 3 was enriched for B cell-B cell and B cell-T cell contacts and depleted of contacts with neutrophils and macrophages, the organization expected of lymphoid aggregates. Niche 4 showed the complementary pattern, with enriched neutrophil-neutrophil and neutrophil-macrophage contacts and few lymphoid contacts. The two tumor niches differed in the same way as their compositions: niche 1 was depleted of nearly all immune contacts, whereas niche 2 retained contacts with endothelial and immune cells, consistent with its peripheral location.

Because the two immune niches carry opposite signs, their relative abundance defines a single prognostic axis per patient. Stratifying all 407 patients into tertiles of a two-niche score, the z-scored log share of niche 3 minus that of niche 4, separated overall survival over 15 years of follow-up (three-group log-rank p = 0.020; Fig. 5e). Niche 3-dominant patients (n = 136) had the best survival and niche 4-dominant patients (n = 88) the worst, with 15%, 21% and 34% of patients dying within 3 years in the niche 3-dominant, intermediate (n = 183) and niche 4-dominant groups, and the score was also associated with lower mortality when analyzed as a continuous variable (Cox hazard ratio 0.89 per unit, p = 0.003).

The polarity of the LUAD niches matches the survival analysis of the source study. Sorin et al. reported B-cell frequency as the cell type most strongly associated with longer overall survival, and found that the survival advantage of B-cell-enriched neighborhoods was greatest when those neighborhoods were co-enriched for CD4+ helper T cells, which is the composition of niche 3 [53]. Conversely, neutrophils and CD163+ macrophages were enriched in high-grade solid tumors, HIF1α+ neutrophils were associated with worse overall survival, and neutrophils interacted preferentially with regulatory T cells in high-grade tumors [53], consistent with the adverse neutrophil–macrophage niche 4 and with the neutrophil-rich relapse niches recovered in the NSCLC cohort. GRASS-MIL therefore recovers, from cell-type labels and spatial geometry alone, the lymphoid-myeloid axis that the original study identified through marker-level analyses, and it identifies a concordant direction of association in two independent lung cohorts.

## 3 Discussion

Spatial tissue analyses commonly characterize which cells or niches are present. GRASS-MIL complements such analyses by identifying which local spatial arrangements are predictive of clinical phenotypes, where they are located, and how they relate to one another. It answers this at three linked scales: the model produces a patient-level prediction, decomposes that prediction into signed neighborhood-level contributions, and organizes the contributing neighborhoods into recurrent tissue niches that can be interpreted with conventional spatial statistics. Prediction and interpretation are therefore coupled in the same output: not only a phenotype score, but a spatial readout of the local architectures on which that score depends.

GRASS-MIL differs from existing spatial-niche methods primarily in this coupling between prediction and interpretation. On the controlled benchmark, GRASS-MIL and SPACE-GM [28] both reached ceiling performance, consistent with clean, fully learnable synthetic signals. On the clinical cohorts, GRASS-MIL achieved the highest mean AUROC among the evaluated methods, exceeding SPACE-GM, CytoCommunity [32], SpaCEy [33] and the composition baselines (Figs. 3g, 4h and 5f). The principal distinction lies in interpretation. CytoCommunity discovers tissue cellular neighborhoods but does not assign them signed contributions to a phenotype prediction. SPACE-GM does decompose its prediction, because the sample-level score is the mean of per-subgraph predictions, but the decomposition is uniform: every neighborhood counts equally, so the local readout is the neighborhood’s own prediction rather than a measure of how much the sample-level decision relies on it. SpaCEy adds a post-hoc explainer that scores neighborhoods by unsigned importance, locating what the model relies on without indicating the direction of the effect. GRASS-MIL produces, within the same model that makes the prediction, an attribution that is signed, weighted by learned relevance, and exact.

The first interpretive layer is the attribution, which follows directly from how GRASS-MIL makes its predictions. The model assigns each neighborhood an attention weight, which acts as an importance weight, and a class-specific prediction score, and the product of the two is the neighborhood’s contribution to the patient-level prediction. These contributions add up exactly to the prediction, so each neighborhood carries a signed contribution: it either moves the prediction toward one class or toward the other. The calculation is performed after training, but it is not a separate explanation method that approximates the model’s behavior; it is the prediction itself, separated into its parts. The distinction matters because attention weights alone are not necessarily reliable indicators of what a model relies on [54]. GRASS-MIL therefore does not treat high attention as evidence of importance on its own, but combines it with the neighborhood’s class-specific score to determine whether, and in which direction, that neighborhood moves the overall prediction.

The side-by-side display of empirical association and attribution in Figs. 3b, 4b and 5a makes a point that applies beyond any one cohort: the two do not always agree, and the disagreement is informative. Empirical association is a marginal statistic, the fraction of a niche’s cells drawn from positive-class regions, weighted by cell count and therefore by section size. Attribution is conditional, the contribution of a niche to a decision made with every other neighborhood of the sample in view. In HGSC, tumor-dominated niches whose cells came almost entirely from untreated sections received no attribution, consistent with the stromal niches already carrying the discriminative information; in NSCLC, the collagen-CAF niche had the highest relapse fraction of any niche but a modest margin, with the adjacent neutrophil-rich niches carrying the signal; in LUAD, the two measures agreed. A niche can therefore be strongly enriched in one class without being what the model relies on, and ranking niches by class fraction would misidentify the structures that matter for prediction. This is the practical reason an exact, signed, within-sample decomposition is needed rather than a post-hoc tally of niche prevalence.

The second interpretive layer is the geometry of the learned neighborhood space. Across the synthetic benchmark and the three clinical cohorts, the learned representations exhibited a recurring star-shaped geometry, with dense recurrent niches at the periphery and intermediate neighborhoods closer to the center. This organization resembles the geometry considered in archetypal analysis, where observations are represented relative to a set of extremal configurations [37, 38]. Related structures have been described in gene-expression spaces, in which extremal states correspond to specialized transcriptional programs and intermediate states combine features of multiple programs, including along spatial axes of a tissue [55, 56, 57]. We interpret the dense HDBSCAN niches analogously as recurrent neighborhood configurations, while the Background population contains heterogeneous or transitional neighborhoods that are not robustly assigned to a dense mode. HDBSCAN clustering turns this geometry into discrete niches for downstream analysis, while its Background label absorbs transitional or impure neighborhoods that do not belong robustly to any dense niche. This combination of continuous geometry, density-based niche definition and explicit residual assignment is central to the biological readout: it lets the model summarize local evidence without forcing every neighborhood into an apparently interpretable category.

In HGSC, the main result is methodological: a single model trained across CosMx/SMI, Xenium and MERFISH data, harmonized only at the level of cell-type labels, recovered a treatment-associated stromal axis whose two architectures correspond to the fibro-inflammatory and desmoplastic stromal changes reported after neoadjuvant chemotherapy [43, 44]. That this signal resides in stromal organization rather than in malignant-cell state, and survives reduction to a ten-type vocabulary, suggests that cell-type maps may suffice for a portable, platform-independent readout of treatment exposure.

In NSCLC, GRASS-MIL linked relapse prediction to a compartmentalized immune-myeloid architecture. The niches associated with favorable outcome were adaptive-immune and TLS-like, in line with reports associating intratumoral lymphoid structures with favorable NSCLC outcomes [52]; the relapse-associated niches were myeloid- and granulocyte-rich, in line with studies linking neutrophil-rich inflammatory states to recurrence and poor outcome [50, 51], with the caveat that neutrophil associations in NSCLC can depend on histology [49]. Spatial statistics added a layer that composition alone does not provide: the two groups of niches occupied mutually exclusive tissue territories rather than intermingling.

The NSCLC cohort also illustrates how GRASS-MIL relates to the analysis that produced the data. The relapse-associated territory contained a CAF niche composed of the collagen, hypoxic, tumor-like and matrix CAF phenotypes that Cords et al. associated with poor prognosis and immune exclusion [45]; the model thus recapitulated the cohort’s established CAF biology from cell-type labels and geometry alone. Within the same territory, however, the strongest attribution fell on neutrophil- and myeloid-rich neighborhoods adjacent to, yet compositionally and spatially distinct from, the CAF niche. Where the original CAF-centered analysis registered neutrophils only as a compositional correlate of the poor-prognosis patient groups [45], GRASS-MIL identifies the myeloid neighborhoods themselves as the dominant contributors to the relapse-associated margin. The arrangement of these niches suggests a hypothesis: matrix-dense stroma borders hypoxic tumor tissue occupied by neutrophils, attribution peaks at the neutrophil end of this sequence, and hypoxic tumor tissue without neutrophils contributes nothing. This pattern is consistent with a spatial link between stromal organization, hypoxia and neutrophil-rich microenvironments and relapse, and with the LUAD finding that the HIF1α+ neutrophil subset, but not total neutrophil frequency, was prognostic [53]. The present data establish the arrangement, not the direction or mechanism of these relationships. Testing them would require the marker intensities excluded from the model, such as neutrophil activation markers where a panel includes them, together with longitudinal or perturbational data.

In LUAD, the same lymphoid-versus-myeloid polarity reappeared in an independent lung cohort with a different endpoint. A B-cell and CD4+ T-cell niche was protective and a neutrophil and classical-macrophage niche adverse for 3-year mortality, whereas the two tumor niches carried no attribution, so the prognostic signal resided in the immune compartment and its contact structure rather than in tumor content. Because both lung cohorts place neutrophil-rich neighborhoods on the adverse side and lymphoid aggregates on the protective side, the concordant direction of association across two independently profiled lung cohorts suggests that the pattern is not specific to a single cohort, panel or annotation scheme. The LUAD analysis also shows that niche-level attribution can be used to derive a patient-level score: the difference in log share between the favorable- and adverse-outcome-associated niches stratified overall survival, which suggests a route from signed neighborhood attribution to simple, interpretable risk indices.

These biological findings are hypothesis-generating rather than causal. GRASS-MIL identifies spatial arrangements that are used by a trained predictive model, not tissue structures that have been experimentally shown to drive treatment response or relapse. Moving from predictive architecture to mechanism will require orthogonal validation, perturbational evidence, or prospective cohorts. The distinction is especially important for attribution: the neighborhood contributions are exact with respect to the trained model, but they inherit whatever the model has learned. An exact explanation of a biased or confounded decision is still an explanation of that decision, not proof of biological causality.

Several limitations follow from the design. First, portability comes at the cost of molecular resolution. Representing each cell by type, optionally with size and inter-cell distance, allows one architecture to span incompatible panels, but it discards marker intensity and continuous cell state. Phenotypes encoded in expression gradients rather than in categorical tissue organization may therefore not be captured by the current model. Second, niche definitions depend on the trained embedding and on density-clustering parameters. Finally, the spatial-statistic readouts are descriptive. They characterize the architectures associated with model predictions but do not test whether those architectures are functionally required.

A natural next step is to combine GRASS-MIL’s exact MIL decomposition with richer, transferable spatial representations. Spatial foundation models are beginning to address the tension between continuous molecular detail and cross-assay portability, including transcriptomic and proteomic settings with explicit panel- or protocol-harmonization mechanisms [24, 25, 26, 27]. Such models could replace or augment the current cell-type-based neighborhood encoder, while preserving the gated-attention decomposition and niche-level spatial readout introduced here. In that sense, GRASS-MIL is a step toward spatial models that not only predict from tissue, but identify the local architectures on which those predictions depend.

## 4 Methods

### 4.1 Datasets and cohorts

We evaluated GRASS-MIL on one controlled simulation benchmark and three published spatial single-cell cancer cohorts: a high-grade serous tubo-ovarian carcinoma (HGSC) cohort from Yeh et al. [41], a non-small-cell lung cancer (NSCLC) imaging mass cytometry cohort from Cords et al. [45, 46] and a lung adenocarcinoma (LUAD) imaging mass cytometry cohort from Sorin et al. [53]. The simulation benchmark was used to test recovery of known spatial signals under controlled generative mechanisms. The three clinical cohorts were used to evaluate phenotype prediction and downstream niche interpretation in real spatial tissue data.

#### Controlled simulation benchmark

The simulation benchmark was designed to separate global composition, local composition and graph-encoded spatial arrangement. Synthetic tissue cores were generated on a square domain containing rectangular or elliptical compartments. Cells were sampled from a uniform Poisson process, assigned one of five cell types according to the simulated compartment structure and connected by Delaunay triangulation. Phenotype generators varied three spatial axes: mixing degree, boundary tortuosity and focal-structure presence. These mechanisms produced ten binary phenotype tasks spanning global mixing, boundary geometry, density and embedded-niche settings. For each simulated core, the same cellular graph construction and ego-graph extraction pipeline was used as for the clinical cohorts (described below). Boundary-proximal ego-graphs were excluded from evaluation to avoid finite-window artifacts in Delaunay-derived ego-graph topology. In particular, ego-graphs centered near the edge of the simulated domain have truncated neighborhoods because part of their *k*-hop context lies outside the sampled window. The simulation benchmark was therefore evaluated on interior ego-graphs. The ego-graph radius *k* was selected by phenotype-specific ablation, following the same procedure used for the clinical cohorts (below); cohort- and task-specific values are reported in Supplementary Table 1a.

#### HGSC spatial transcriptomics cohort

The HGSC cohort was obtained from Yeh et al. [41], who generated a single-cell spatial transcriptomic atlas of high-grade serous tubo-ovarian carcinoma. The atlas contains 130 tumors from 94 patients and more than 2.5 million high-quality spatially resolved single-cell measurements across CosMx/SMI, Xenium and MERFISH platforms. We used the processed cell-level data, cell centroids and cell-type annotations from the source atlas. The prediction task was treatment-status classification, defined as neoadjuvant-chemotherapy-treated versus treatment-naive at the time of tissue collection. Because segmentation-derived cell-size measurements were not consistently available across all platforms, the HGSC model used cell-type identity as the only node feature. Cell centroids were used to construct cellular graphs and edge-distance features. Gene-expression values, marker intensities and platform-specific molecular measurements were not used as model inputs. Model performance was estimated by nested 5-fold cross-validation over the 196 tissue regions (folds of 44, 34, 36, 43 and 39 regions). Regions were grouped so that all regions from one patient, including pre- and post-treatment biopsies, fell into the same fold, and folds were stratified on treatment status crossed with acquisition platform so that the label was balanced across folds. All HGSC performance metrics were computed on the outer test folds.

#### NSCLC imaging mass cytometry cohort

The NSCLC cohort was obtained from Cords et al. [45, 46], who profiled tissue microarrays by imaging mass cytometry (IMC). The dataset comprises 1,070 patients and 2,070 tissue microarray cores, with one or two cores available per patient. The IMC panel contained 45 markers and was designed to resolve tumor, stromal, immune and cancer-associated fibroblast (CAF) states in the NSCLC tumor microenvironment. We used the processed single-cell table, cell centroids, cell-type annotations and segmentation-derived cell-size measurements from the source dataset. The prediction task was relapse classification. We retained patients annotated as adenocarcinoma, squamous-cell carcinoma or adenosquamous carcinoma and excluded other histologic categories. Cell-type labels were taken from the original publication without modification. For GRASS-MIL input features, each cell was represented by its annotated cell type and segmentation area. Cell centroids were used to construct cellular graphs and edge-distance features. Marker intensities were not used as model inputs. Model performance was estimated by nested 5-fold cross-validation over the 1,868 cores retained after histology filtering (folds of 375, 372, 372, 375 and 374 cores). Cores were grouped into patients and folds were defined with StratifiedGroupKFold (random seed 0), stratified on the patient-level relapse label with every core of a patient confined to one fold. Model training and inference were performed at the core level, treating each tissue microarray core as a separate tissue region, and NSCLC performance was reported at the core level on the outer test folds.

#### LUAD imaging mass cytometry cohort

The LUAD cohort was obtained from Sorin et al. [53], who profiled tissue microarrays from 416 patients with lung adenocarcinoma by IMC. One 1-mm^2^ core was available per patient, acquired with a 35-marker panel at approximately 1 µm resolution. The published annotation resolves tumor cells, endothelial cells, 14 immune populations and an undefined class, giving the 17 cell types used here. We used the processed single-cell table, cell centroids and cell-type annotations from the source dataset. The prediction task was death within 3 years of surgery versus known survival beyond 3 years; patients censored before 3 years were excluded, leaving 407 labeled patients with 89 deaths. For GRASS-MIL input features, each cell was represented by its annotated cell type and segmentation area. Cell centroids were used to construct cellular graphs and edge-distance features, and marker intensities were not used as model inputs. Because each core corresponds to one patient, cross-validation folds were defined with StratifiedKFold over the 416 patients (folds of 84, 83, 83, 83 and 83 patients), stratified jointly on age above 75 years, sex and death; after exclusion of censored patients the folds contained 83, 82, 82, 81 and 79 labeled patients. All LUAD performance metrics were computed on the outer test folds.

#### Cell-type handling and filtering

For all three clinical cohorts, cell-type annotations were taken from the processed source publications without modification. Cells without valid centroid coordinates or cell-type labels were removed before graph construction.

#### Ethics and data availability

All clinical data were obtained from previously published studies. Ethical approval, informed consent procedures and primary data-generation protocols are described in the original publications [41, 45, 53]. This study used de-identified processed data and did not involve additional human-subject sample collection.

### 4.2 Cellular graph construction and ego-graph sampling

For each tissue region, we constructed an undirected cellular graph *G* = (*V, E*) whose nodes correspond to segmented cells and whose edges encode local spatial adjacency. Node positions were the two-dimensional centroids of segmentation masks. Edges were defined by Voronoi-polygon adjacency: two cells were connected when their Voronoi polygons shared a boundary, equivalently when they were adjacent in the Delaunay triangulation of the centroid point set. Each node *v* ∈ *V* carried a categorical cell-type label *c* _*v*_ ∈ {1, …, *T*} and, where available, a continuous cell-size feature *s*_*v*_ given by the segmentation area. The set of model-visible node features was cohort-specific. In the NSCLC and LUAD cohorts, the model used cell type and cell size. In the HGSC cohort, where cell size was not available consistently across platforms, the model used cell type only. No marker intensities, gene-expression values or biomarker measurements were used as model inputs. Each edge (*u, v*) ∈ *E* carried a single continuous feature: the Euclidean centroid distance *d*_*uv*_ . The natural-log distance was clipped to the interval [2, 5] and linearly rescaled to [0, 1], identically for all cohorts, before being passed to the graph encoder.

For each anchor cell *v*, we extracted the induced *k*-hop graph centered on the cell (ego-graph) containing all cells reachable from *v* within *k* message-passing steps. An additional Euclidean radius cutoff excluded spatially distant cells connected through long triangulation edges. The ego-graph radius *k* was selected by validation or simulation-specific ablation rather than fixed globally; most reported models used *k* = 4, while some simulation settings selected *k* = 3. Each ego-graph was treated as one MIL instance, and the collection of ego-graphs from a tissue region formed the corresponding MIL bag. Cohort- and experiment-specific values of *k* and associated radius cutoffs are reported in Supplementary Table 1a.

### 4.3 Graph encoder

Each ego-graph was encoded with a graph neural network based on GINEConv layers [34, 58]. Cell-type labels were mapped to learned embeddings. Continuous node features, when present, were linearly projected to the same dimension and added to the cell-type embedding. Edge distances were encoded with a learned continuous edge-feature projection and used during message passing. At layer *l*, node representations were updated as

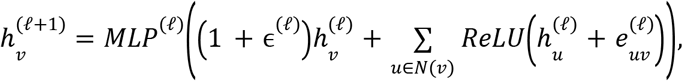

where 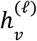 is the node representation at layer *l*, 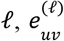 is the edge-distance embedding for edge (*u, v*), and ϵ^(*l*)^ is a trainable scalar. The encoder used four message-passing layers with hidden dimension 256. Layer normalization, ReLU activations and dropout were applied between layers. A graph-level max pool over the final node representations produced one ego-graph embedding

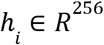

for each instance *i*. Cohort-specific hyperparameters, including feature availability, selected *k*, dropout, learning rates, batch sizes and sampling parameters, are reported in Supplementary Table 1.

### 4.4 Self-supervised encoder pretraining

Before supervised training, the graph encoder was pretrained using Bootstrapped Graph Latents (BGRL) [35]. For each input ego-graph, two independently augmented graph views were generated. We used the graph augmentations selected on the basis of the evaluation in [59]: importance-based node dropping, importance-based edge dropping and feature perturbation. Node and edge importance scores were computed from local graph structure, so that augmentations preferentially perturb less structurally informative components while preserving the principal local graph structure. Feature perturbations were applied to continuous node features when such features were available. For the HGSC cohort, which used cell-type labels only, the continuous-feature perturbation step was omitted. BGRL uses an online encoder *f*_θ_ and a target encoder *f*_ξ_ with the same architecture. The target encoder parameters were updated as an exponential moving average of the online encoder parameters,

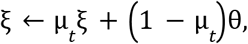

with µ_*t*_ following a cosine schedule from 0.99 to 1.0. A predictor MLP was applied to the online representation. Writing

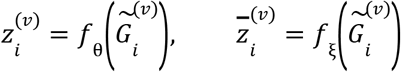

for view *v* ∈ {1, 2} of ego-graph *i*, the pretraining objective was the symmetric negative cosine similarity

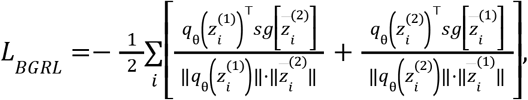

where *q*_θ_ is the online predictor and *sg*[·] denotes stop-gradient. Pretraining hyperparameters are reported in Supplementary Table 1b.

### 4.5 Attention-based multiple-instance prediction head

Each tissue region *r* was represented as a bag *B* of ego-graph instances. The graph encoder produced an embedding h_*i*_ ∈ *R* ^256^ for each instance *i* ∈ *B*_*r*_. These embeddings were pooled with a gated-attention MIL head [14] (Fig. 1a). The encoder output was first projected to a lower-dimensional representation,

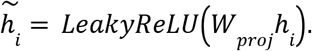

The gated-attention module then computed

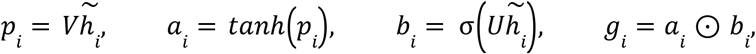

where *p* is the pre-tanh activation of one gate, σ is the elementwise logistic sigmoid and ⊙ denotes the Hadamard product. The pre-tanh space 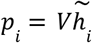 was used as the primary representation space for embedding-geometry analysis and niche discovery. A class-wise linear map produced per-instance attention logits

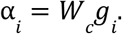

Attention weights were normalized over instances within each bag and separately for each class,

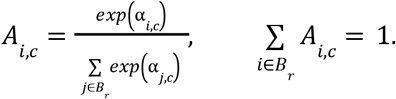

In parallel, a per-instance MLP mapped each encoder embedding h_*i*_ to class logits *l*_*i,c*_ . The bag-level class logit was the attention-weighted sum of instance logits,

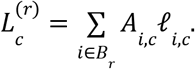

Class probabilities were obtained by applying the softmax over the C class logits; all tasks in this study are binary (C = 2), so the predicted positive-class probability is the softmax over two logits. Models were trained with region-level, class-weighted cross-entropy, with class weights set from the inverse class frequencies of the training split (HGSC and NSCLC) or, for LUAD, binary cross-entropy with a positive-class weight,

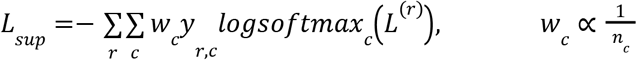

Optimization used AdamW [60] with cosine learning-rate decay. Model selection used validation AUROC: within the nested cross-validation, the epoch with the highest inner-cross-validation AUROC was retained for the refit, and the LUAD model stopped training when validation AUROC had not improved for 10 epochs. Hyperparameters were selected on validation data and are listed in Supplementary Table 1b.

### 4.6 Exact neighborhood attribution

GRASS-MIL attribution follows directly from the additive form of the MIL prediction head. For each instance *i* and class *c*, we define the class-specific contribution

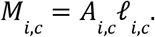

The bag-level class logit is exactly the sum of these instance contributions,

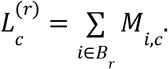

This decomposition holds for any number of classes C, and any linear contrast of the class logits inherits it. For a one-versus-rest comparison of class c against the remaining classes, we define the class-c margin contribution of instance i as its contribution to class c minus the average of its contributions to the other classes,

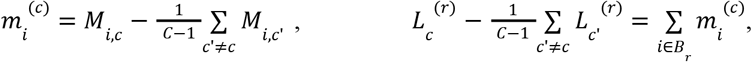

so that the one-versus-rest bag margin 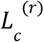 decomposes exactly. (A contrast against the maximum of the other classes would not decompose, because a maximum of sums is not a sum of maxima.) All tasks in this study are binary (C = 2), for which this reduces to the signed class-1-versus-class-0 margin contribution

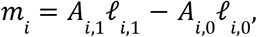

so that the bag margin decomposes as

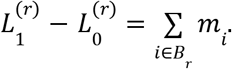

This identity is exact for the trained model and does not require a baseline, perturbation, path integral or separately fitted explainer. Because attention weights are normalized over the bag, individual contributions scale with the number of instances in the region. We therefore report the bag-size-scaled contribution

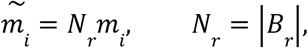

when comparing per-neighborhood contributions across regions. For niche-level attribution, the statistic reported in the figures as the relative-attention margin is the mean of the size-scaled contributions over the instances assigned to niche *k* in region *r*,

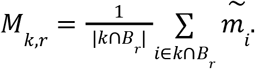

For this calculation, the bias of the final layer of the per-instance classifier was omitted from the instance logits, so that contributions reflect instance-level evidence rather than the class prior. The niche-level summary is the unweighted mean of these per-region averages over the regions in which the niche occurs,

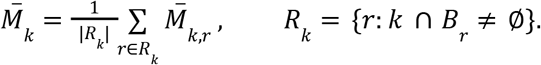

This statistic weights every region equally, irrespective of how confident the model is about it, and summarizes whether a niche contributes positively or negatively to the corresponding decision; uncertainty was estimated by bootstrap over regions (below).

### 4.7 Embedding geometry and niche definition

We analyzed the representation space given by the pre-tanh gated-attention activation

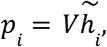

where 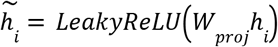 is the projected GNN embedding. This is the representation entering one gate of the gated-attention head before the tanh nonlinearity. It was used for both geometric visualization and niche discovery. As a model-independent baseline, we also computed a local composition vector for each ego-graph, defined as the *L*_1_ -normalized cell-type count vector over the nodes of the ego-graph. The pre-tanh representations were reduced by principal component analysis, and the first two principal components were used to visualize the large-scale geometry of the learned neighborhood space. Recurrent tissue niches were defined using HDBSCAN [36] in cosine space. HDBSCAN cluster-size and related parameters are reported in Supplementary Table 1a. Points assigned by HDBSCAN to the noise label were retained as a separate unassigned Background population. We interpret this Background population as neighborhoods that do not belong robustly to any high-density niche, including heterogeneous, transitional or weakly structured neighborhoods. Star-shaped geometry was assessed qualitatively in projections onto the first principal components. In this geometry, dense HDBSCAN niches occupy the arms of the embedding space, while intermediate neighborhoods lie along transitions between them through its center. We interpret these dense niches as analogous to the extremal configurations represented by vertices in archetypal analysis [37, 38, 56], under the assumption that recurrent neighborhood configurations are populated densely enough to form modes; neighborhoods that are not are assigned to Background rather than forced onto a vertex, and the polytope itself is not fitted. Niche-level attribution and spatial statistics were computed using the HDBSCAN assignments.

### 4.8 Spatial characterization of niches

Spatial statistics were used to characterize the cellular organization of the niches identified in the pre-tanh representation space, covering cell-type composition, Moran’s *I*, Ripley’s *L* and cross-*L*, permutation-based neighborhood enrichment and, for LUAD, within-niche contact enrichment (below). Unless otherwise noted, statistics were computed per tissue region and then aggregated across regions.

#### Cell-type composition

For each niche, cell-type composition was computed as the mean cell-type frequency across all ego-graphs assigned to that niche. Composition profiles were used to annotate niches and to compare learned spatial motifs with local cell-type abundance.

#### Moran’s *I*

Spatial autocorrelation of cell-type indicators was quantified with global Moran’s *I* [39]. For a cell-type indicator *x*_*v*_ = 1[*c*_*v*_ = *t*] on a graph with adjacency weights *w*_*vu*_, Moran’s *I* was

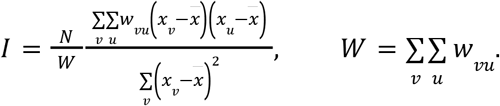

Binary graph adjacency was used for *w*_*vu*_. Moran’s *I* was computed for selected cell types within each niche to quantify whether those cell types were spatially coherent or dispersed. Significance was assessed by label permutation with Benjamini–Hochberg correction across tested niche and cell-type pairs.

#### Ripley’s *L* and cross-*L*

Second-order spatial structure was quantified with Ripley’s *K* and *L* functions [40]. For a point pattern of intensity λ in a region of area |*A*|,

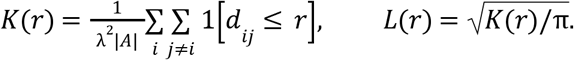

We report the centered form *L*(*r*) − *r*, which is zero under complete spatial randomness. For pairs of cell types or niche-assigned cell populations, we computed the bivariate cross-function

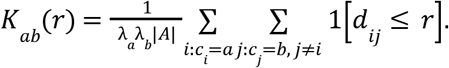

Positive centered cross-*L* values indicate co-aggregation above random expectation, whereas negative values indicate spatial exclusion. For summary heatmaps, we evaluated centered cross-*L* at the characteristic radius *r*^*^ that maximized the mean absolute centered cross-*L* across pairs.

#### Neighborhood enrichment

Cell-type-pair neighborhood enrichment was computed using permutation-based enrichment analysis as implemented in `squidpy` [9]. Cell-type labels were permuted over the fixed spatial graph to estimate enrichment relative to random labeling. The resulting *z*-scores were used to identify overrepresented and underrepresented cell-type adjacencies within each niche.

#### Within-niche contact enrichment

For the LUAD cohort, cell-type-pair organization within niches was summarized by a contact-frequency ratio rather than by permutation. For each niche and each ordered pair of cell types, the frequency of contacts between the two types among all cell contacts inside the niche was divided by the frequency of the same contacts among all contacts in the whole tissue, and the ratio was reported on a log2 scale, so that positive values indicate contacts enriched inside the niche and negative values contacts depleted. Pairs with fewer than 100 contacts inside the niche were masked.

### 4.9 Model evaluation and statistical analysis

Model performance on the clinical cohorts was evaluated by nested 5-fold cross-validation. Within each outer fold, hyperparameters and early-stopping points were selected by AUROC on inner cross-validation or on a held-out validation fold drawn from the training folds, and never on the outer test fold. The primary metric was the area under the receiver operating characteristic curve (AUROC) on the outer test fold, reported as the mean ± s.d. across the five outer folds. For the simulation benchmark, AUROC was computed separately for each phenotype generator and summarized by the mean and worst-case AUROC across generators. Baselines included global cell-type composition classifiers, local neighborhood-composition classifiers, and the spatial graph-based baselines described above (SPACE-GM [28], CytoCommunity [32] and SpaCEy [33]). All baselines were run on the same harmonized cell-type input as GRASS-MIL, with cell size where available and no marker intensities, to keep the input classes comparable across methods; SPACE-GM and SpaCEy were additionally trained on their native marker-intensity inputs, and those results are reported in Supplementary Fig. 1 and Supplementary Table 3. Global composition baselines used whole-region cell-type frequency vectors as input. Local composition baselines used cell-type frequency vectors computed on the same ego-graphs as GRASS-MIL and aggregated predictions to the region level. Full baseline definitions, model families, hyperparameters and cohort-specific evaluation details are provided in Supplementary Table 2, and the fold-level AUROC of every model on every clinical endpoint in Supplementary Table 3. Spatial-statistic significance was assessed with permutation tests and Benjamini–Hochberg false-discovery-rate correction within each figure.

### 4.10 Implementation

The pipeline was implemented in Python using PyTorch [61] and PyTorch Geometric [62]. Spatial analyses used `squidpy` [9], `scanpy` [63], and `scikit-learn` [64]. Code for GRASS-MIL is available at https://github.com/BoevaLab/grass-mil.

