## Supplementary Information (Supplementary Figures 1-3 and Supplementary Tables 1-3) for "GRASS-MIL: graph-based representation and discovery of phenotype-associated spatial structures with multiple instance learning"

#### Supplementary Figures

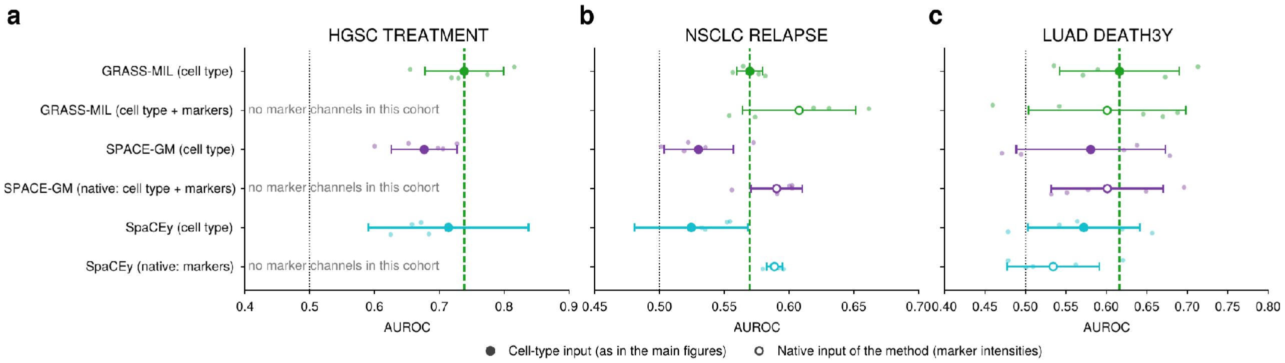


**Supplementary Figure 1 | Spatial baselines with their native marker-intensity inputs.**

Test AUROC of the three spatial-niche methods on (**a**) HGSC treatment status, (**b**) NSCLC relapse and (**c**) LUAD 3-year death, in the format of Figs. 3g, 4h and 5f: large markers, mean over the five outer test folds of the nested 5-fold cross-validation on the shared, patient-grouped partition; horizontal bars, s.d. across folds; small dots, individual folds; dotted line, chance; dashed green line, mean AUROC of GRASS-MIL with cell-type input. Filled markers denote the cell-type input used in the main figures (one-hot cell type, plus segmentation area in NSCLC and LUAD). Open markers denote the method's native input: for SPACE-GM the original node features (cell type, area and per-cell marker intensities), for SpaCEy per-cell marker intensities only, and for GRASS-MIL the same cell-type-plus-marker features as SPACE-GM (encoder pretrained and fine-tuned on that input). Marker intensities are the 43 IMC channels of the NSCLC cohort and 19 IMC channels of the LUAD cohort (per-cell mean intensity over the segmentation mask); the HGSC spatial-transcriptomics cohort has no marker channels, so only cell-type rows are shown. CytoCommunity is defined on cell-type labels only and therefore has no native-marker variant. Numeric performance values are listed in Supplementary Table 3.

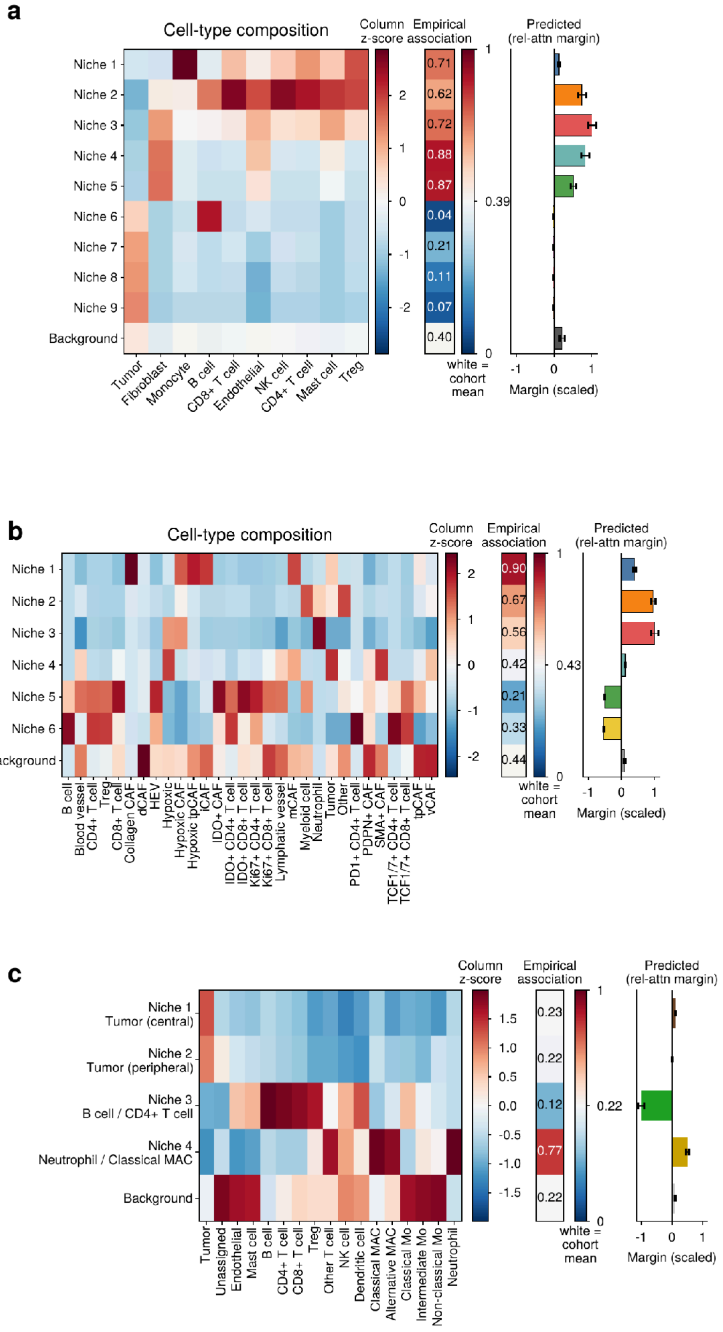


**Supplementary Figure 2 | Niche cell-type composition as column-wise z-scores.**

The composition heatmaps of Fig. 3b (**a**, HGSC, niches 1–9 and Background × 10 cell types), Fig. 4b (**b**, NSCLC, niches 1–6 and Background × 30 cell types) and Fig. 5a (**c**, LUAD, niches 1–4 and Background × 17 cell types), with each cell-type column z-scored across the niche rows (red, cell type over-represented in the niche relative to the other niches; blue, under-represented). The underlying values are the same mean per-ego-graph cell-type fractions shown as percentages in the main figures; z-scoring exposes cell types that are rare in absolute terms (for example the CAF subtypes in NSCLC niche 1 and the hypoxic tumor cells and neutrophils in NSCLC niches 3 and 4). Empirical association, the fraction of cells in the niche that come from positive-class regions, with the diverging color map centered on the cohort mean (white; 0.39 HGSC, 0.43 NSCLC, 0.22 LUAD). Predicted, the relative-attention-weighted margin contribution of the niche (Methods), scaled so that the largest absolute niche margin in each cohort equals 1; bars are colored by niche, error bars are 95% bootstrap confidence intervals over regions.

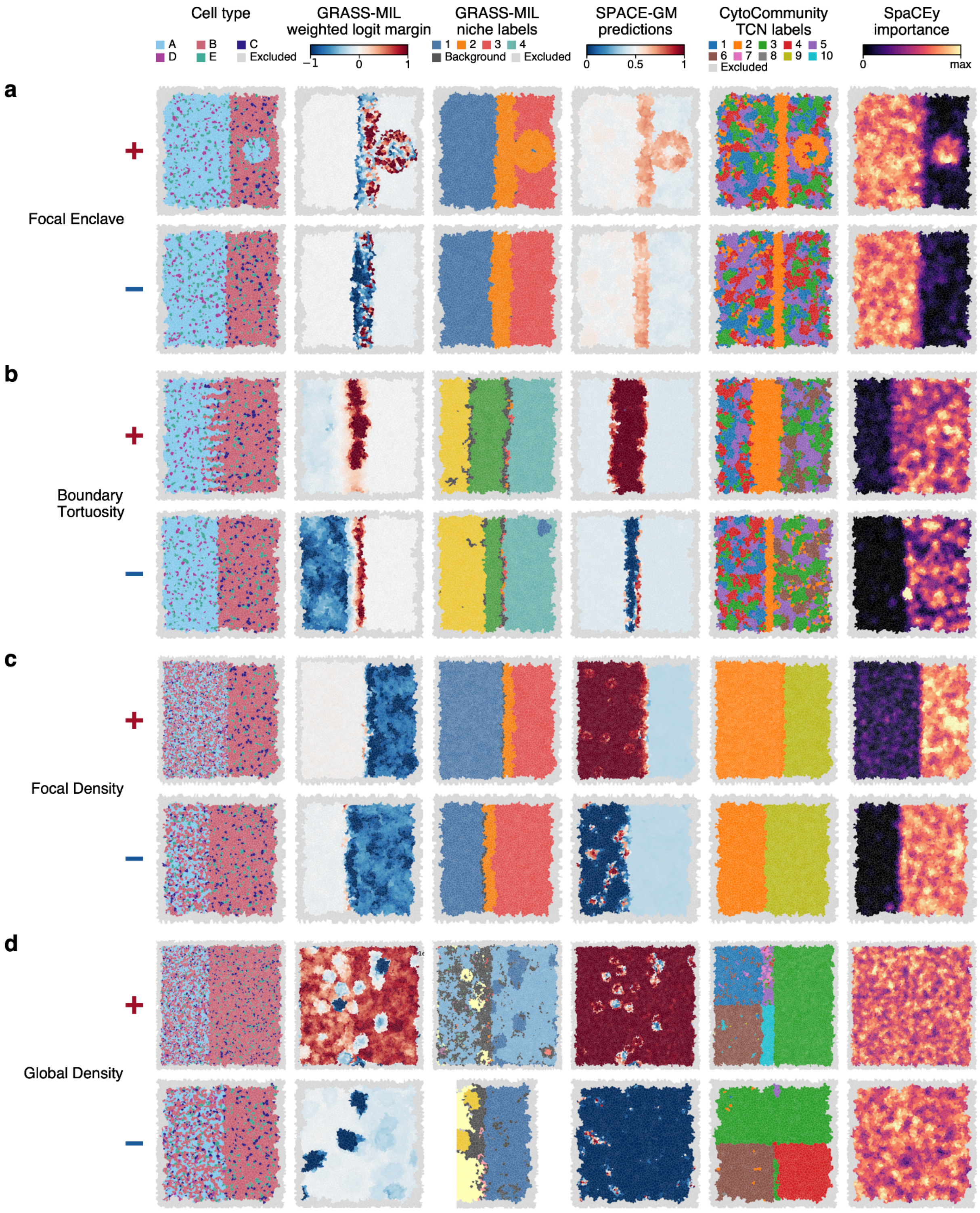
Cell type
A
B
C
D
E
Excluded
GRASS-MIL weighted logit margin
−1
0
1
GRASS-MIL niche labels
1
2
3
4
Background
Excluded
SPACE-GM predictions
0
0.5
1
CytoCommunity TCN labels
1
2
3
4
5
6
7
8
9
10
Excluded
SpaCEy importance
0
max
a
+
−
Focal Enclave
b
Boundary Tortuosity
c
Focal Density
d
Global Density

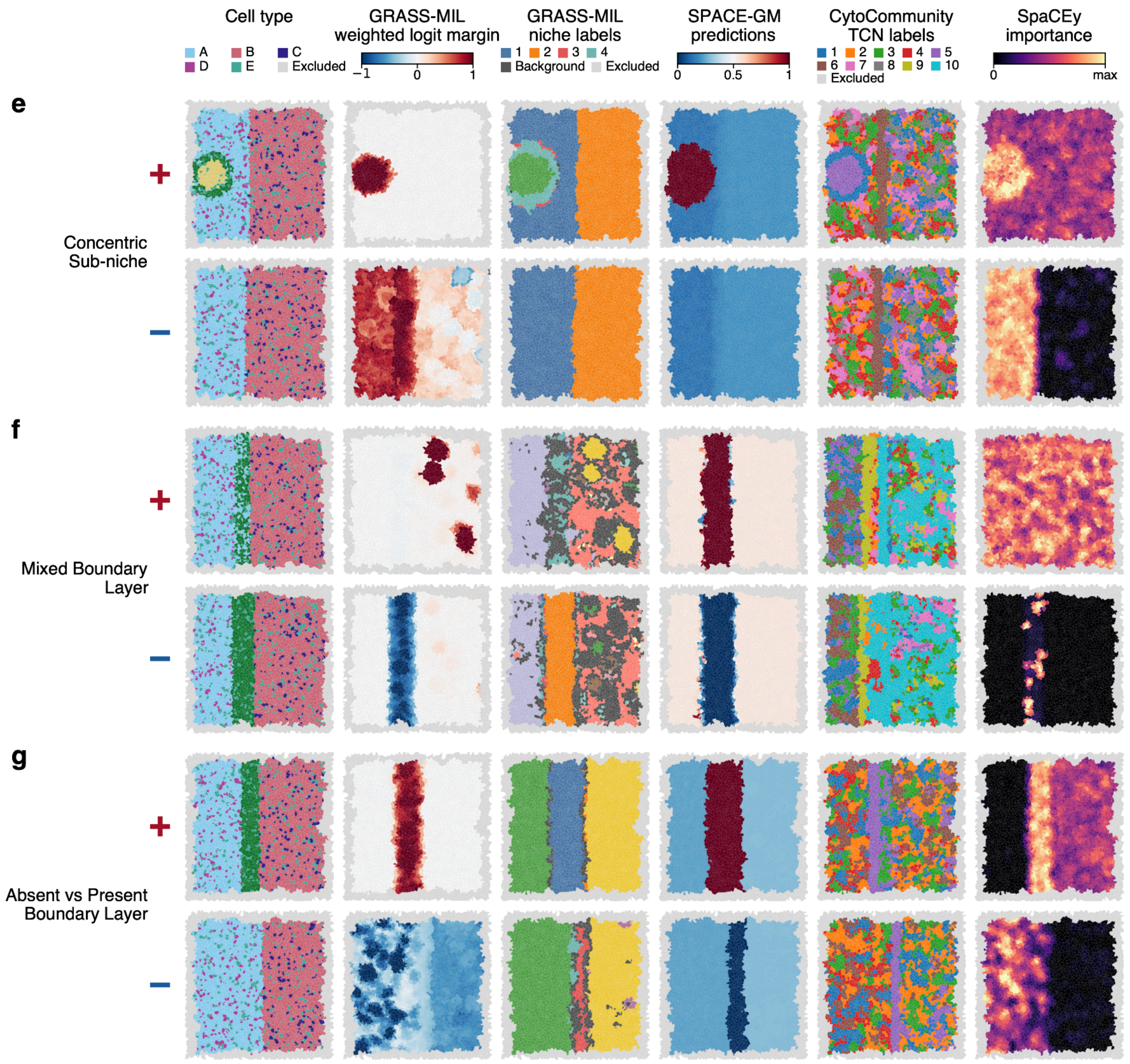


**Supplementary Figure 3 | Example cores for the simulation generators not shown in Fig. 2d–f.**

One positive (+) and one negative (−) test core for each of the seven remaining generators, rendered as in Fig. 2d–f: (a) Focal Enclave, (b) Boundary Tortuosity, (c) Focal Density, (d) Global Density, (e) Concentric Sub-niche, (f) Mixed Boundary Layer, (g) Absent vs Present Boundary Layer. Columns, left to right: simulated cell type (five types, A–E); GRASS-MIL bag-size-scaled margin contribution of the ego-graph centered on each cell (red, towards the positive class; blue, towards the negative class); GRASS-MIL niche label of each ego-graph (HDBSCAN on the pre-tanh attention space, Background = unassigned); SPACE-GM per-ego-graph predicted probability of the positive class; CytoCommunity tissue cellular neighborhood (TCN) label of each cell; SpaCEy node importance (GNNExplainer edge masks, top-20 % threshold, mean over the 2-hop neighborhood, scaled from 0 to the 99th percentile of the core). Cells in the 4-hop boundary rim are excluded from evaluation and shown in grey. Example cores are the ones used in the simulation evaluation report (the positive and negative test cores with the median bag score of each class). The cores shown are held-out test cores.

### Supplementary Tables

**Supplementary Table 1 | Cohort- and task-specific configuration.**

Data, cellular-graph construction, evaluation protocol and niche-discovery settings (**a**) and encoder pretraining, MIL-head training and model-selection settings (**b**) for the simulation benchmark and the three clinical cohorts. Ego-graph radius k and the Euclidean radius cutoff (75k + 55 coordinate units) apply to graph construction, pretraining, training and the local-composition baselines alike. HDBSCAN is run on L2-normalized principal components (cosine geometry). BGRL, Bootstrapped Graph Latents; EMA, exponential moving average; eom, excess-of-mass cluster selection.

#### 1a. Data, cellular graph, evaluation and niche discovery

| Setting | Simulation (10 generators) | HGSC TREATMENT | NSCLC RELAPSE | LUAD DEATH3Y |
|---|---|---|---|---|
| Regions (patients) | 900 synthetic cores; each generator labels a subset (fixed split 615 / 180 / 105 train / val / test regions; 82 / 24 / 14 labeled per generator) | 196 regions, 94 patients, 5 datasets, 3 platforms | 1,868 TMA cores (ADC, SCC, adenosquamous), ~1,000 patients | 416 cores = 416 patients; 407 labeled for DEATH3Y (89 deaths) |
| Positive class (prevalence) | generator-specific (balanced) | post-treatment biopsy (0.39 of cells) | relapse (0.43 of cells) | death ≤ 3 y (0.22 of cells) |
| Cell types (node vocabulary) | 5 types (A–E) | 11 (10 annotated + Other) | 30 | 17 |
| Node features available | cell type | cell type | cell type, segmentation area | cell type, segmentation area |
| Node features used by the model | cell type (one-hot → learned embedding) | cell type | cell type + area (area ÷ cohort 95th-percentile area) | cell type + area (area ÷ cohort 95th percentile) |
| Marker intensities | none | Not used (variable) | not used (43 channels; native-input variant only, Supplementary Fig. 1) | not used (19 channels; native-input variant only) |
| Graph edges | Delaunay triangulation (Voronoi adjacency) | Delaunay triangulation (Voronoi adjacency) | Delaunay triangulation (Voronoi adjacency) | Delaunay triangulation (Voronoi adjacency) |
| Edge feature | continuous: clip((ln d − 2) / 3, 0, 1), d = centroid distance in coordinate units (GINEConv) | continuous: clip((ln d − 2) / 3, 0, 1), d = centroid distance in coordinate units (GINEConv) | continuous: clip((ln d − 2) / 3, 0, 1), d = centroid distance in coordinate units (GINEConv) | continuous: clip((ln d − 2) / 3, 0, 1), d = centroid distance in coordinate units (GINEConv) |
| Ego-graph radius k / Euclidean cutoff | k = 4; 75k + 55 = 280 units | k = 4; 355 units | k = 4; 355 units | k = 4; 355 units |
| Boundary handling | evaluation and niche analysis restricted to interior cells (4-hop rim excluded) | evaluation and niche analysis restricted to interior cells (4-hop rim excluded) | evaluation and niche analysis restricted to interior cells (4-hop rim excluded) | evaluation and niche analysis restricted to interior cells (4-hop rim excluded) |
| Outer evaluation | fixed train / val / test split, test AUROC per generator | nested 5-fold CV; folds 44/34/36/43/39; grouped by clinical fingerprint (patient), stratified on treatment × platform | nested 5-fold CV; folds 375/372/372/375/374; StratifiedGroupKFold (seed 0) grouped by patient fingerprint (survival tuple + TMA block), stratified on relapse | nested 5-fold CV; folds 84/83/83/83/83 (83/82/82/81/79 labeled); StratifiedKFold (seed 0) stratified on age ≥ 75 × sex × death |
| Inner selection | validation split (val AUROC / val loss) | inner 3-fold CV on the development folds | inner 3-fold CV on the development folds | inner 3-fold CV on the development folds |
| Niche space | pre-tanh attention activation | pre-tanh attention activation | pre-tanh attention activation | pre-tanh attention activation |
| PCA before clustering | 10 PCs | 10 PCs | 8 PCs (visualization 2 PCs) | 10 PCs |
| HDBSCAN | min_cluster_size 100, cluster_selection eom, euclidean on L2-normalized PCs (cosine) | min_cluster_size 100, cluster_selection leaf, L2-normalized PCs → 9 niches + Background | min_cluster_size 300, eom, L2-normalized PCs → 6 niches + Background | min_cluster_size 150, eom, L2-normalized PCs → 4 niches + Background |
| Cells / ego-graphs clustered | test regions (interior), sample 200,000 | held-out test regions, sample 20,000 | 50,000 sampled held-out ego-graphs | 79 fold-4 test cores, sample 50,000 |

#### 1b. Training and model selection

| Setting | Simulation | HGSC TREATMENT | NSCLC RELAPSE | LUAD DEATH3Y |
|---|---|---|---|---|
| Encoder | GIN, 3 layers, width 256, GIN-MLP width 512, max pool, LayerNorm | GIN, 4 layers, width 256, max pool; LayerNorm | GIN, 4 layers, width 256, max pool, LayerNorm | GIN, 4 layers, width 256, max pool, LayerNorm |
| BGRL pretraining: iterations, batch | 50,000 it., 64 ego-graphs | 200,000 it., 64 | 200,000 it., 64 | 200,000 it., 64 |
| BGRL: optimizer | AdamW, lr $5 \times 10^{-5}$, weight decay $10^{-5}$, 1,000 warm-up steps, cosine decay | AdamW, lr $5 \times 10^{-5}$, weight decay $10^{-5}$, 1,000 warm-up steps, cosine decay | AdamW, lr $5 \times 10^{-5}$, weight decay $10^{-5}$, 1,000 warm-up steps, cosine decay | AdamW, lr $5 \times 10^{-5}$, weight decay $10^{-5}$, 1,000 warm-up steps, cosine decay |
| BGRL: EMA momentum | 0.99 - 1.0, cosine schedule | 0.99 - 1.0, cosine schedule | 0.99 - 1.0, cosine schedule | 0.99 - 1.0, cosine schedule |
| BGRL: predictor | MLP, hidden 512 | MLP, hidden 512 | MLP, hidden 512 | MLP, hidden 512 |
| BGRL: augmentations (both views) | DropImportance (degree-based importance; μ = 0.25, p_λ = 0.45): cell-type masking to the unassigned type and importance-weighted edge dropping; no feature noise (no area feature) | DropImportance (μ = 0.25, p_λ = 0.45); no feature noise (no area feature) | DropImportance (μ = 0.25, p_λ = 0.45) + Gaussian noise on the area feature (σ = 0.2) | DropImportance (μ = 0.25, p_λ = 0.45) + Gaussian noise on the area feature (σ = 0.2) |
| Encoder dropout in pretraining | 0 | 0.1 | 0.1 | 0.1 |
| Pretraining regions | all 900 cores | all 196 regions (label-free) | all 1,868 cores (label-free) | all 416 cores (label-free) |
| MIL head: attention | gated attention, projection p ∈ {16, 32, 64}, D ∈ {4, 8, 16} (swept) | projection 64, D 8 | projection 32, D 8 | projection 32, D 8 |
| Instance head | 3-layer MLP (256-256-2) | 3-layer MLP (256-256-2) | 3-layer MLP (256-256-2) | 3-layer MLP (256-64-1) |
| Loss | cross-entropy (bag level) | class-weighted cross-entropy (inverse train-split frequency) | class-weighted cross-entropy (inverse train-split frequency) | BCE with positive-class weight (1 − p) / p, p = train-fold positive rate |
| Model / epoch selection | checkpoint with best validation macro-AUROC inside each run | epoch with the highest inner-CV validation AUROC (3 inner folds), then refit on all development folds for that epoch with early stopping disabled | epoch with the highest inner-CV validation AUROC (3 inner folds), then refit on all development folds for that epoch with early stopping disabled | epoch with the highest inner-CV validation AUROC (3 inner folds), then refit on all development folds with early stopping disabled |

**Supplementary Table 2 | Baseline definitions.**

(**a**) The three spatial-niche methods: implementation, the input used in the main-figure comparison (cell type, plus segmentation area where available), the native input used in Supplementary Fig. 1, architecture and training recipe, the hyper-parameter search and the selection rule applied inside each outer fold, and cohort-specific deviations. CytoCommunity was run in its supervised (phenotype-labeled) mode; its tissue cellular neighborhoods (TCNs) are learned transductively on all regions, and only the downstream abundance classifier and the number of TCNs are cross-validated. (**b**) The five classifier families used as composition baselines, in the global setting (one whole-region cell-type proportion vector per region) and the local setting (one 4-hop ego-graph proportion vector per cell, region score = mean cell-level output), with their feature definitions and search spaces for the clinical cohorts and for the simulation benchmark. All clinical baselines share the nested 5-fold protocol and the fixed partitions of Supplementary Table 1a.

All clinical-cohort baselines are evaluated on the same fixed, patient-grouped 5-fold partition as GRASS-MIL (Supplementary Table 1a); every hyper-parameter or checkpoint choice is made inside the four development folds of each outer fold, never on the test fold. Simulation baselines use the fixed train / val / test split of the benchmark.

#### 2a. Spatial-niche methods

| Method | Implementation | Input used here (main figures) | Native input (Supplementary Fig. 1) | Architecture / training | Hyper-parameter search and selection | Cohort-specific notes |
|---|---|---|---|---|---|---|
| SPACE-GM (Wu et al.) | `spacegm` package, reference configuration of the published `run_gin.py` | one-hot cell type (+ area in NSCLC / LUAD), categorical neighbor / distant edge type (55-unit cutoff) | cell type + area + per-cell marker intensities (NSCLC: 43 IMC channels; LUAD: 19 IMC channels) | 3-hop ego-graphs; randomly initialized GIN, 3 layers, embedding 512, BatchNorm, max pool, dropout 0.25; MLP head; class-weighted cross-entropy; Adam lr $10^{-3}$; 50,000 iterations of 64 ego-graphs; region score = mean P(positive) over 96 sampled ego-graphs | no search (reference configuration); checkpoint evaluated every 2,500 iterations on the validation fold (outer fold i + 1), best-validation-AUROC checkpoint kept; train = remaining 3 folds | Simulation: 3-hop GIN, embedding 512, max pool, 3-layer MLP head, Adam lr $10^{-3}$, 50,000 iterations, per-generator model, best validation checkpoint |
| CytoCommunity (Hu et al.; CytoCommunity-plus) | authors' Step1–Step4 pipeline (soft TCN assignment, supervised mode) | cell-type labels + coordinates (the method's only input) | — (cell-type only by design) | Step 1: tissue split into patches (quadtree, ≤ 1,000 cells per patch); Step 2: k-NN cell graphs (k from the mean patch size); Step 3: graph-level supervised TCN learning (GraphLabel = region phenotype, 2 classes, MinCut pooling), 10 runs × 200 epochs, mini-batch 16, Num_TCN ∈ {8, 10, 12}; Step 4: consensus TCN assignment over runs; per-region TCN-abundance vector (fraction of cells per TCN) | prediction: balanced logistic regression (C = 1) on the standardized TCN-abundance vector, trained on the development folds; Num_TCN chosen per outer fold by inner 3-fold CV | TCNs are learned once per cohort on all regions with their labels (transductive, as the method is designed), so the TCN features have seen the test labels; this favors the baseline. Simulation: Num_TCN ∈ {6, 8, 10}, 10 runs × 400 epochs, k-NN k = 20, selection by validation AUROC |
| SpaCEy (Saez lab) | authors' `bin/model.py` (CustomGCN, GATv2 variant), re-run under the shared CV | one-hot cell type (+ area in NSCLC / LUAD) on the authors' Delaunay graph | per-cell marker intensities only (the method's native formulation) | Delaunay graph with per-image pruning of edges above the 97.5 % length quantile; 2 GATv2 layers (1 head) + 2 feed-forward layers (width 128), dropout 0.2, mean pool, single logit, BCE-with-logits; Adam, weight decay $10^{-5}$, ReduceLROnPlateau (factor 0.5, patience 10, min lr $10^{-4}$); batch 16 graphs; ≤ 200 epochs; early stopping on validation loss (patience 20) with checkpoint kept when loss and AUROC both improve | grid over hidden width {32, 64} × lr {$10^{-3}$, $10^{-2}$}; per outer fold the validation fold (outer fold i + 1) selects the configuration and checkpoint by validation AUROC; train = remaining 3 folds | Simulation: 48 configurations per non-trivially-solved generator sampled from the authors' HPO space (hidden {16, 32, 64, 128}, FF width {64, 128, 256}, dropout {0.2, 0.3}, lr {$10^{-4}$, $10^{-3}$, $10^{-2}$}, weight decay {$10^{-5}$, $10^{-4}$, $10^{-3}$}, 2–3 GCN layers, 1–2 FF layers, 1–2 heads, batch {16, 32}), selected by validation AUROC |

#### 2b. Composition classifiers

Five classifier families, each in a *global* and a *local* setting.

| Setting | Feature per training example | Prediction for a region | Regions / examples |
|---|---|---|---|
| Global | region cell-type proportion vector (11 / 30 / 17 types; standardized on the training folds) | classifier output | one example per region |

| | | | |
|---|---|---|---|
| Local | 4-hop ego-graph cell-type proportion vector of a single cell (same k as GRASS-MIL, graph adjacency without the Euclidean cutoff); region label broadcast to its cells | mean of the classifier output over all cells of the region | 300 cells sampled per development region (200 in NSCLC) so that regions are weighted equally |

| **Classifier** | **Clinical cohorts (global and local)** | **Simulation (global and local, scikit-learn defaults unless stated)** |
|---|---|---|
| LogReg | balanced L2 logistic regression; C ∈ {$10^{-5}$, …, 10} by inner 3-fold CV (LogisticRegressionCV, AUROC) | L2 logistic regression, C = 1 |
| RandomForest | 500 trees, min leaf 5, balanced class weights (local: 300 trees, min leaf 20) | 100 trees |
| SVM | RBF kernel, balanced class weights; C ∈ {0.1, 1, 10, 100} × γ ∈ {scale, 0.01, 0.1, 1} by inner 3-fold grid search, scored by decision function (local: C = 1, γ = scale on a 15,000-cell subsample) | RBF kernel, C = 1, γ = scale, probability outputs |
| GradBoost | histogram gradient boosting, depth 3, learning rate 0.05, 300 rounds, L2 = 1, balanced sample weights | gradient boosting, 100 rounds, depth 3 |
| MLP | one hidden layer of 64, α = $10^{-2}$, early stopping, positives oversampled to balance | two hidden layers of 512, adaptive learning rate, early stopping |

Local baselines in the simulation sample ≤ 64 interior cells per region as ego-graph centres (same 4-hop radius) and average the per-cell probabilities.

**Supplementary Table 3 | Test AUROC of every model on every clinical endpoint.**

Mean ± s.d. of the AUROC over the five outer test folds of the nested 5-fold cross-validation. All rows within a cohort use the same fixed, patient-grouped partition and are therefore paired. Rows 1-14 use cell-type input (one-hot cell type; plus segmentation area for the graph models in NSCLC and LUAD) and are the rows plotted in Figs. 3g, 4h and 5f; the last three rows are the native-marker variants plotted in Supplementary Fig. 1 (not available for HGSC, which has no marker channels). "local" denotes the instance-level composition learners (one training example per cell, its 4-hop ego-graph cell-type composition, region score = mean of the cell-level outputs); unsuffixed classifier rows are trained on whole-region composition vectors. The nested-CV protocol of every baseline is given in Supplementary Table 2.

| Model | Input | HGSC TREATMENT | NSCLC RELAPSE | LUAD DEATH3Y |
|---|---|---|---|---|
| GRASS-MIL | cell type | 0.739 ± 0.061 | 0.570 ± 0.010 | 0.616 ± 0.074 |
| SPACE-GM | cell type | 0.677 ± 0.051 | 0.530 ± 0.027 | 0.581 ± 0.092 |
| CytoCommunity | cell type | 0.601 ± 0.061 | 0.522 ± 0.009 | 0.493 ± 0.073 |
| SpaCEy | cell type | 0.714 ± 0.124 | 0.525 ± 0.044 | 0.572 ± 0.069 |
| LogReg local | cell type | 0.720 ± 0.120 | 0.542 ± 0.008 | 0.590 ± 0.066 |
| RandomForest local | cell type | 0.683 ± 0.073 | 0.548 ± 0.006 | 0.594 ± 0.059 |
| SVM local | cell type | 0.678 ± 0.097 | 0.533 ± 0.008 | 0.594 ± 0.029 |
| GradBoost local | cell type | 0.680 ± 0.065 | 0.555 ± 0.005 | 0.588 ± 0.043 |
| MLP local | cell type | 0.688 ± 0.098 | 0.541 ± 0.007 | 0.593 ± 0.048 |
| LogReg | cell type | 0.688 ± 0.089 | 0.549 ± 0.012 | 0.592 ± 0.060 |
| RandomForest | cell type | 0.626 ± 0.072 | 0.553 ± 0.010 | 0.515 ± 0.072 |
| SVM | cell type | 0.658 ± 0.093 | 0.519 ± 0.008 | 0.572 ± 0.059 |
| GradBoost | cell type | 0.616 ± 0.085 | 0.545 ± 0.011 | 0.487 ± 0.094 |
| MLP | cell type | 0.657 ± 0.108 | 0.538 ± 0.008 | 0.571 ± 0.085 |
| GRASS-MIL | cell type + markers | n/a | 0.608 ± 0.044 | 0.601 ± 0.097 |
| SPACE-GM | native: cell type + markers | n/a | 0.590 ± 0.020 | 0.601 ± 0.069 |
| SpaCEy | native: markers | n/a | 0.589 ± 0.006 | 0.534 ± 0.057 |
